# Ethylene modulates cell wall mechanics for root responses to compaction

**DOI:** 10.1101/2025.03.02.640043

**Authors:** Jiao Zhang, Zhuo Qu, Zengyu Liu, Jingbin Li, Edward Farrar, Osvaldo Chara, Lucas Peralta Ogorek, Augusto Borges, Shingo Sakamoto, Nobutaka Mitsuda, Xiaobo Zhu, Mingyuan Zhu, Jin Shi, Wanqi Liang, Malcolm Bennett, Bipin Pandey, Dabing Zhang, Staffan Persson

## Abstract

Soil stresses impact crop yields, presenting global agricultural challenges. Soil compaction triggers root length reduction and radial expansion driven by the plant hormone ethylene. We report how ethylene controls cell wall properties to promote root radial expansion. We demonstrate how soil compaction stress, via ethylene, upregulates *Auxin Response Factor1* in the root cortex, which represses Cellulose Synthase (*CESA*) genes. *CESA* repression drives radial expansion of root cortical cells by modifying the thickness and mechanics of their cell walls, which result in a “stiff epidermis-soft cortex” contrast. Our research thus connects ethylene signaling with root mechanics via cell wall strength, and reveals how dynamic regulation of cellulose synthesis crucially controls root growth in compacted soil.

## Introduction

Modern agriculture’s reliance on mechanization is causing soil degradation and compaction, impacting root growth and crop yield^1^. Plant roots radially expand when encountering compacted soil conditions, leading to shorter and thicker roots^2^. Radial root swelling is mainly due to expansion of cortex cell layers^3^ that can cause soil fissures that aid soil penetration^2,4^. This adaptive response is driven by entrapment of the plant volatile hormone ethylene around the root as compaction reduces gas diffusion through the soil^5,6^. Despite this mechanistic insight, how ethylene controls cell wall properties to enable root radial expansion remains unclear.

Primary plant cell walls, and in particular the organization and amount of the load-bearing polymer cellulose, support anisotropic cell expansion and are key to organ growth^7^. Cellulose is produced at the plasma membrane by large, multimeric protein complexes known as Cellulose Synthase (CESA) complexes (CSCs)^8^. In rice, OsCESA3, 5, 6, and 8 form heterotrimeric CSCs that are active in growing root cells during primary wall synthesis^9^. When roots encounter soil compaction, the arrangement of cellulose microfibrils in their cortical cell walls is altered^10^. While the mechanical strength of cell walls is crucial for anisotropic cell expansion, little is known about how cell wall properties regulate root penetration of soil.

### Reduced cellulose synthase activity promotes root penetration of compacted soil

To explore whether changes in cellulose synthesis impact the ability of roots to penetrate compacted media, we initially grew rice seedlings for five days on normal (0.3%) and dense (0.6%) agar media, with and without varying concentrations of Indaziflam. Indaziflam is a cellulose biosynthesis inhibitor (CBI) that suppresses cellulose production by triggering an atypical increase in CSC density at the plasma membrane in plant cells^11, 12^. Surprisingly, while high concentrations (250 pM) of the inhibitor suppressed root growth (Extended Data Fig. 1a,b), low levels (150 pM) promoted root penetration through the dense growth media (Extended Data Fig. 1a,b). To validate our results in soil conditions, we adopted a genetic approach and mutated *OsCESA6* using CRISPR-Cas9 technology. *OsCESA6* encodes a redundant CESA subunit (Extended Data Fig. 2a) that can be replaced by other CESAs in the CSCs^13^ and therefore exhibits mild cellulose developmental defects when mutated^14^. We identified a *cesa6* mutant line that carried a deletion of an ‘A’ in the fourth exon causing early termination of the gene (Extended Data Fig. 2b). Non-invasive X-ray CT imaging revealed that *cesa6* mutant roots grew longer than wild type (WT) in compacted (1.6 g/cm^3^) than non-compacted (1.2 g/cm^3^) soil conditions, as well as on dense agar media (Fig. 1a,b; Extended Data Fig. 2c,d), corroborating our Indaziflam results. Notably, while *cesa6* mutant plants typically have shorter and swollen roots compared to wild type when grown on media plates, the mutant and wild-type roots were of equal lengths when grown in germination pouches (Fig. 1c,d). These results indicate that minor perturbations in cellulose biosynthesis impact the ability of plant roots to extend under varying soil and media conditions.

**Fig. 1:**
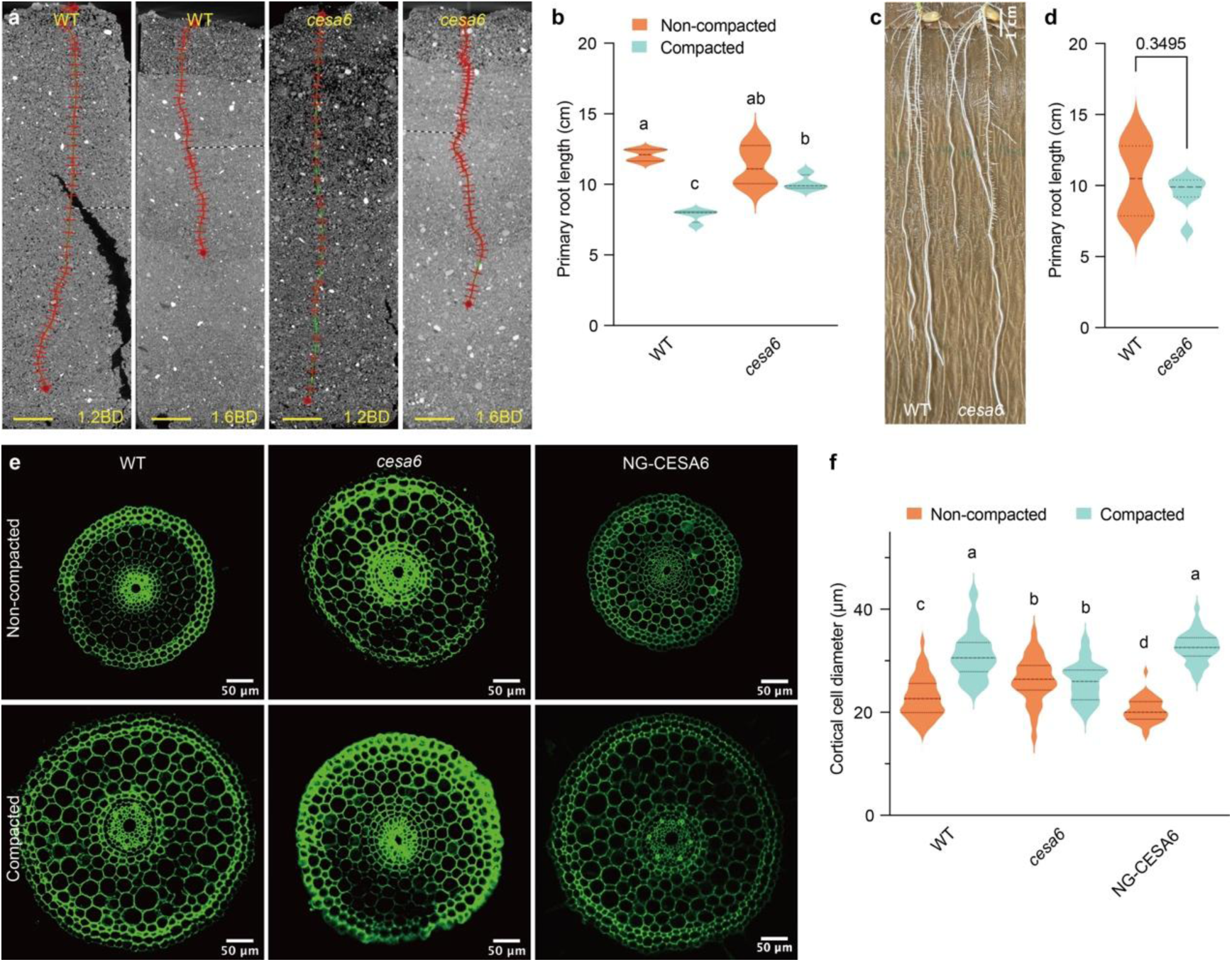
OsCESA6 negatively regulates rice root penetration in compacted soil. **a-b,** Primary root growth of 5-day-old WT and *cesa6* mutants cultivated in non- compacted (1.2 g cm⁻³ bulk density [BD]) and compacted (1.6 g cm⁻³ BD) soil conditions. The images shown are representative samples from 4 independent CT scans. Statistical analysis was conducted using two-way ANOVA and Tukey test, with different letters indicating significant differences at the 0.05 level, *p* (interaction) = 0.0032. Scale bars in (a), 10 mm. **c-d,** Primary root growth of 7-day-old wild-type (WT) and *cesa6* mutants grown in the germination bags. The experiment was replicated three times, consistently yielding similar trends (n=8 for each trial). Statistical analysis was performed using Welch’s t-test with a significance threshold of 0.05, based on data from one representative trial. **e,** Root cross-sections of 5-day-old WT, *cesa6*, and NG-CESA6 (*cesa6* complementation lines) from the elongation zones. Plants were grown in normal (0.3%) and dense (0.6%) media conditions. Sections were prepared using a vibratome at 50 µm thickness and observed under UV light at 405 nm using an SP5 confocal microscope. Scale bar: 50 µm. **f,** Cortical cell diameters of WT, *cesa6*, and NG-CESA6, measured from sections as described in (e). Measurements were taken from 40 cells across 5 sections for each genotype. The experiment was replicated three times, consistently yielding similar trends. Statistical analysis was performed using two-way ANOVA and Tukey test, with different letters indicating significant differences at the 0.05 level, *p* (interaction) < 0.0001, based on data from one representative trial.

We next examined root cross-sections of *cesa6* and Indaziflam-grown seedlings and found that they exhibited larger cortical cell diameter than WT in non-compacted conditions, but similar dimensions in compacted conditions (Fig. 1e,f; Extended Data Fig. 1c,d). These growth phenotypes were restored to WT in the *cesa6* mutant by complementation using a mNeonGreen-tagged CESA6 construct driven by its native promoter (NG-CESA6; Fig. 1e,f; Extended Data Fig. 2c,d)^15^.

### *OsARF1* is upregulated by compaction stress and represses *CESA* gene expression

To establish a connection between the changes in cellulose synthesis and upstream signals such as ethylene, we screened for potential regulators of cellulose synthesis. Genes associated with cellulose synthesis tend to be co-expressed (Extended Data Fig. 3a), indicating that a common set of regulators, i.e. transcription factors (TFs), may control the process similar to that of metabolic regulons^16,17^. Therefore, we conducted a comprehensive yeast-one-hybrid (Y1H) assay using nine promoters from co-expressed core primary wall cellulose synthesis-related rice genes as baits and 1143 rice TFs as preys. By exclusively considering TFs that bind to multiple promoters, we aimed to minimize false positives and enrich “true” TFs regulating primary wall cellulose synthesis. From these assays, we found several ethylene response factors (ERFs), including OsERF34 and 39 (binding to eight and five promoters, respectively; Fig. 2a). These ERFs are the closest homologs of Arabidopsis ERF35^18^, the main primary cell wall regulating TF in Arabidopsis (Extended Data Fig. 3b), thereby confirming the validity of our approach to identify primary cell wall and cellulose synthesis regulators.

**Fig. 2:**
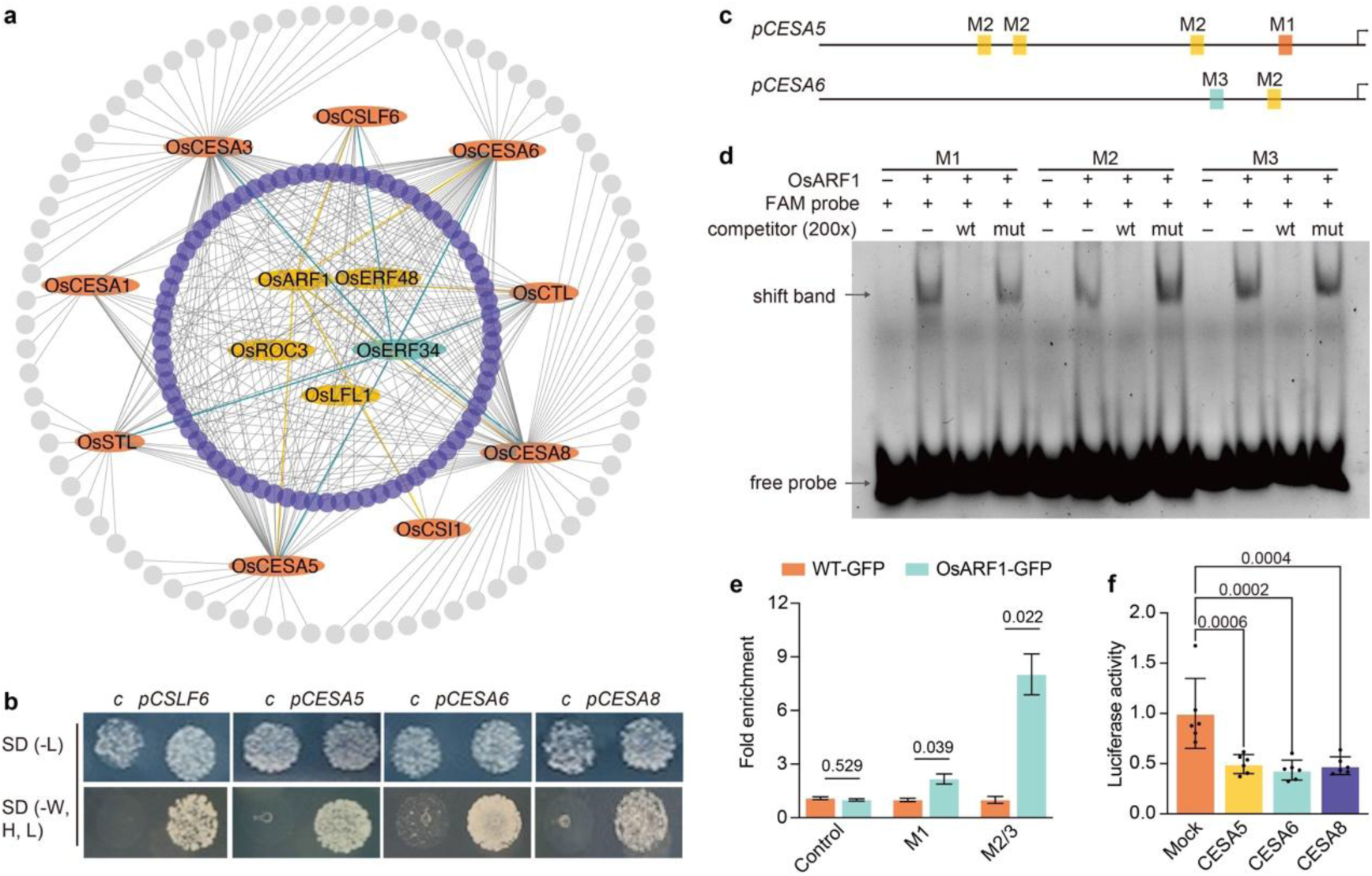
OsARF1 directly binds to several primary wall cellulose-related promoters. **a**, Promoters of *OsCESA1*, *3*, *5*, *6*, *8*, *OsCSLF6*, *OsCSI1*, *OsSTL*, and *OsCTL* (depicted as orange ovals) were used as baits in yeast one-hybrid (Y1H) screens against a library of 1,143 rice transcription factors (TFs). Gray lines represent binding interactions between TFs and baits. The screen identified transcription factors with diverse binding patterns, ranging from highly promiscuous TFs (blue: 8 promoters; yellow: 6 promoters) to those with intermediate (purple: 2-5) or single-promoter (gray) binding specificity. **b**, Y1H assays demonstrate that OsARF1 can directly bind to the promoters of *OsCESA5*, *6*, *8*, and *OsCSLF6*. ‘c’ denotes the empty vector control; SD represents Synthetic Dextrose Minimal Medium; ‘W’ indicates tryptophan; ‘H’ indicates histidine; ‘L’ indicates leucine; and ‘-’ signifies absence. **c**, Analysis of OsARF1 binding cis-elements on the promoters of *OsCESA5* and *6*. Boxes denote AuxRE-L (TGTCNN) cis elements. M1 (orange) represents TGTCACCGACA, M2 (yellow) represents CCGACA, and M3 (blue) represents TGTCAC. **d**, Electrophoretic mobility shift assays (EMSA) evaluating the direct binding of OsARF1 to probes M1-M3 from (c), with competition (200-fold excess of non-labeled probes) and mutated probes (TGTCNN to AAAAAA). Probes were 5’-end labeled with FAM, and interactions were visualized via the Cy2 channel. **e**, Chromatin immunoprecipitation followed by quantitative PCR (ChIP-qPCR) experiments confirm the *in vivo* binding of OsARF1 to M1, M2, M3, and a negative control (a sequence in promoters but without AuxRE-L binding motifs). Results are presented as mean ± SD, based on one of three biological replicates (Welch’s t-test with Bonferroni-Dunn method at 0.05 level). **f**, Transient effector-reporter assays in rice protoplasts showed OsARF1-mediated transcriptional regulation. The experiment was replicated twice, consistently yielding similar trends. In one representative trial, six biologically independent samples were analyzed. Individual data points are represented by dots on the graph (One-way ANOVA with Dunnett test at 0.05 level).

In our Y1H, we also discovered that auxin response factor 1 (OsARF1) bound six of the promoter baits (Fig. 2a). We verified the interactions between OsARF1 and the bait promoters using pairwise Y1H assays, electrophoretic mobility shift assay (EMSA) against potential ARF cis-elements and chromatin immunoprecipitation quantitative PCR (ChIP-qPCR) experiments (Fig. 2b-e). These data corroborated that OsARF1 interacts with several primary wall *CESA* promoters in rice. Next, we analyzed whether OsARF1 could repress or activate the *CESA* promoters using a transient effector-reporter assay system in protoplasts extracted from rice leaves, and found that OsARF1 repressed *CESA* promoter activity (Fig. 2f). Interestingly, *OsARF1* is induced by compacted soil and media conditions (Fig. 3a-c), potentially linking cellulose synthesis regulation and soil compaction. A GUS reporter fusion revealed that *OsARF1* is expressed in the stele when grown in control conditions, but its expression was dramatically induced across other root elongation zone tissues, particularly in the cortex of early elongation zone, in response to dense agar media (Fig. 3a), which we confirmed by RT-qPCR analysis (Fig. 3b). Therefore, we selected *OsARF1* for further analysis.

**Fig. 3:**
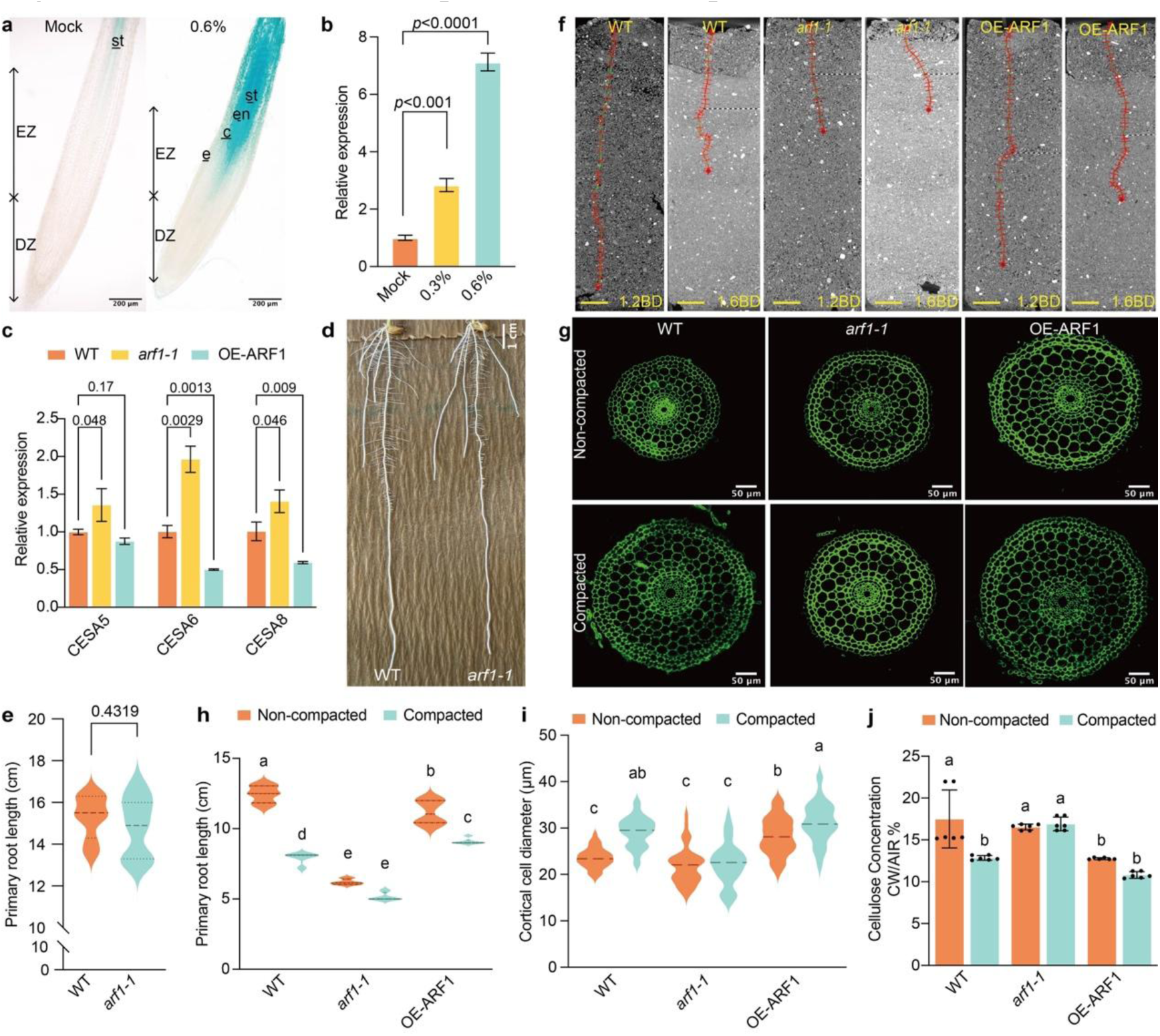
OsARF1 enhances root penetration of compacted soils. **a,** *ProOsARF1*:GUS expression in mock (water) and dense media for 48 hours. Longitudinal sections (50 µm thickness) were prepared using a vibratome. Scale bars: 200 µm. EZ: elongation zone, DZ: division zone, e: epidermis, c: cortex, en: endodermis, st: stele. **b,** *OsARF1* expression in WT root elongation zone under mock (water), normal, and dense media. Statistics presented are from one of three representative trials (One-way ANOVA with Dunnett test at 0.05 level). Bars indicate mean ± SD. **c,** Expression of *CESA5*, *6*, and *8* in WT, *arf1-1*, and OE-ARF1 seedling roots (n=6 per trial). The experiment was performed in triplicate with consistent trends; statistics presented are from one representative trial (One-way ANOVA with Dunnett test at 0.05 level). **d-e,** Primary root length of 9-day-old WT and *arf1-1* grown in the germination bags (n=11 per trial). The experiment was replicated thrice, yielding consistent trends. Statistical analysis was performed using Welch’s t-test (*p* < 0.05) based on one representative trial. **f,** CT scans of primary root length in WT, *arf1-1*, and OE-ARF1 plants grown for 5 days in soil. Root lengths were derived from 4 independent CT images. Scale bar: 1 cm. **g,** Confocal images of root cortical diameter in elongation zones from 5-day-old WT, *arf1-1*, and OE-ARF1 plants grown under normal and dense soil conditions. Scale bar: 50 µm. **h,** Primary root lengths from (f). Statistical analysis was conducted using two-way ANOVA and Tukey test, with different letters indicating significant differences at the 0.05 level, *p* (interaction) < 0.0001. **i,** Cell measurements (40 cells from 5 sections per genotype). The experiment was conducted in triplicate with consistent trends. Data presented are from one representative trial. Statistical analysis was conducted using two-way ANOVA and Tukey test, with different letters indicating significant differences at the 0.05 level, *p* (interaction) = 0.0001. **j,** Cellulose content in WT, *arf1-1*, and OE-ARF1 roots grown under normal and dense conditions for 5 days (n=6). Individual data points are represented by dots. Statistical analysis was conducted using two-way ANOVA and Tukey test, with different letters indicating significant differences at the 0.05 level, *p* (interaction) = 0.001..

### OsARF1 regulates *CESA* expression to control root compaction responses

To investigate OsARF1 function on root growth in compacted soil and media, we generated *osarf1* mutants, namely *arf1-1* and *arf1-2* (Extended Data Fig. 4a), via CRISPR-Cas9 and *OsARF1* overexpression lines (OEs; Extended Data Fig. 5a). We first examined the expression patterns of *CESA* genes in the roots of both the CRISPR-generated mutants and OE lines. The expression levels of *CESA5*, *6*, and *8* were significantly elevated in the *arf1-1* mutant but, with the exception of *CESA5*, suppressed in the OE-ARF1 lines. These findings substantiate that OsARF1 functions as a transcriptional repressor of the *CESA* gene family (Fig. 3c).

To investigate root growth characteristics of *arf1-1* and OE-ARF1, we grew WT, *arf1-1*, and OE-ARF1 lines either on normal and dense agar media, or in non- compacted and compacted soil for five days. WT seedling root length was inhibited by 35% in compacted soil and 24% on dense media (Fig. 3f,h; Extended Data Fig. 4b,c). By contrast, *arf1-1* displayed root growth defects in both non-compacted and compacted conditions (Fig. 3f,h; Extended Data Fig. 4b,c), whereas OE-ARF1 roots were less affected than WT in compacted conditions as roots were inhibited 19% (soil) and 1% (media) (Fig. 3f,h; Extended Data Fig. 5b,c). The *arf1-1* mutant roots were of similar length as wild type when grown in germination bags (Fig. 3d,e), indicating that OsARF1 may be associated with root growth regulation in response to different compaction conditions.

To reveal the effects of OsARF1 activity on the root radial response to compaction, we analyzed root cross-sections from WT, *arf1* mutants, and OE-ARF1. WT roots increased their diameter when subjected to compaction conditions; but *arf1-1* and *arf1- 2* failed to do so (Fig. 3g,i; Extended Data Fig. 4d,e). Intriguingly, OE-ARF1 displayed constitutive root swelling, i.e. in both non-compacted and compacted conditions (Fig. 3g,i), phenocopying *cesa6* mutants and Indaziflam-treated roots. We reasoned that if OsARF1 regulates root radial expansion through OsCESA6, their expression patterns should show opposite trends under compaction. Indeed, using *ProOsCESA6*:GUS reporter lines, we found that *OsCESA6* expression was significantly reduced in the cortex of early elongation zone (Extended Data Fig. 6), precisely where OsARF1 expression was strongly induced by compaction (Fig. 3a).

To validate the regulatory relationship between OsARF1 and cellulose synthesis, we quantified cellulose content in roots grown under varying media conditions. While dense media led to reduced cellulose content in WT roots compared to normal conditions, *arf1-1* maintained similar cellulose levels across both conditions, with significantly higher content than WT under dense media conditions (Fig. 3j). Conversely, OE-ARF1 showed consistently lower cellulose content than WT, particularly under normal conditions (Fig. 3j). Further supporting these results, chemical reduction of cellulose synthesis using either Indaziflam (150 pM) or DCB^19^ (50 nM, which reduces CSC mobility without affecting plasma membrane CSC density) rescued the root growth defects of *arf1-1* mutants (Extended Data Figs. 7-8). Furthermore, the *osarf1* mutant root phenotypes on both normal and dense media conditions were rescued in *arf1-1cesa6* double mutant lines (Extended Data Fig. 9). Collectively, these results demonstrate that OsARF1 orchestrates root responses to compaction through modulation of cellulose biosynthesis by suppressing *CESA* expression.

### Roots adapt their growth to compaction by modifying cortical cell wall thickness

Cell-wall biosynthesis and mechanical properties are fundamental determinants of root elongation and expansion. Although the relationship between wall mechanical properties and cell growth has been extensively studied, certain regulatory mechanisms remain elusive. Previous studies have demonstrated that cell wall thickness and stiffness vary across different root regions and cell types^20^. For instance, cells in the elongation zone exhibit lower stiffness compared to those in the meristem zone, facilitating cell elongation^20^. Furthermore, cells with higher stiffness typically possess thicker cell walls^21,22^. Based on these observations, we hypothesized that radially expanded cortical cells would exhibit thinner and more compliant walls. To test this hypothesis, we first examined cortical cell wall thickness of WT, *arf1-1*, OE-ARF1, and *cesa6* in the root elongation zone. Our TEM (Transmission Electron Microscopy) imaging revealed that the wall thickness of WT cortical cells was reduced in compacted conditions compared to non-compacted conditions (Fig. 4a-c). By contrast, we observed thicker root cortex walls in *arf1-1* in both non-compacted and compacted conditions, whereas OE-ARF1 and *cesa6* maintained thinner walls in both conditions (Fig. 4a-c). Complementary Atomic Force Microscopy (AFM) analysis of cortex cell wall stiffness in the elongation zone (Fig. 4d,e) supported our hypothesis, demonstrating lower stiffness in WT compared to *arf1-1*, with OE-ARF1 and *cesa6* showing the lowest stiffness values (Fig. 4d,e). These findings explain the differential responses to soil compaction: *arf1-1* cortex cells maintained their diameter, while OE- ARF1 and *cesa6* exhibited constitutive radial expansion.

**Fig. 4:**
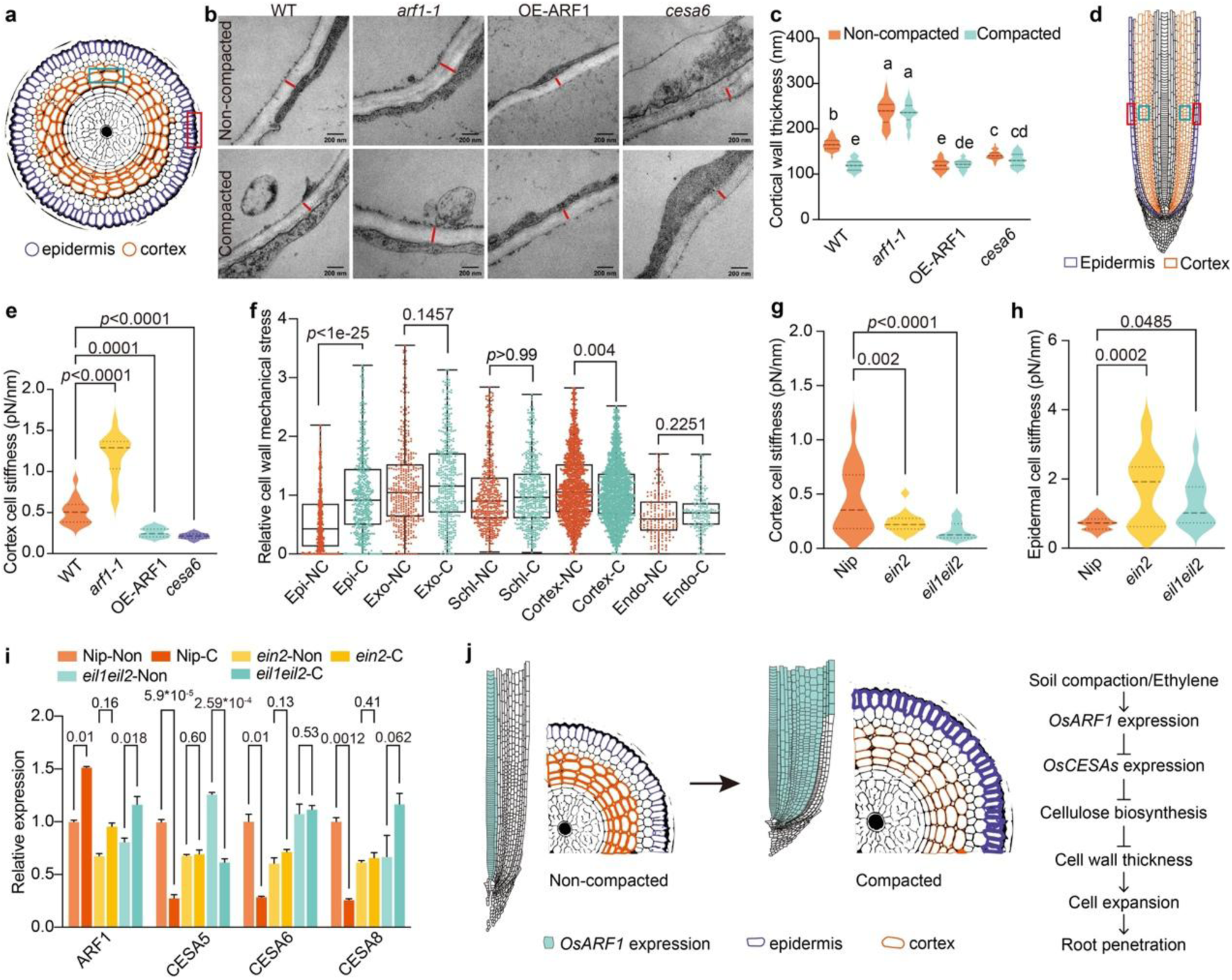
A “stiff-epidermis and soft-cortex” cell wall model facilitates rice root adaption to soil compaction. **a,** Schematic cross-section of a rice root cell, highlighting cortex (blue box) and epidermal (red box) cells. **b-c,** Transmission electron microscopy (TEM) images showing cell wall thickness in cortical cells of 5-day-old WT, *arf1-1*, OE-ARF1, and *cesa6* under normal or dense media (40 cells across 2 sections per genotype). Measurements were repeated twice. Different letters indicate significant differences (two-way ANOVA and Tukey test at 0.05 level, *p* (interaction) < 0.0001). **d,** Longitudinal schematic indicating approximate locations for Atomic Force Microscopy (AFM) force spectroscopy measurements in cortex (blue box) and epidermal (red box) regions. **e,** Cortical cell wall stiffness in WT, *arf1-1*, OE-ARF1, and *cesa6* under dense media conditions. Two sections per genotype were analyzed, with the experiment repeated four times (One-way ANOVA with Dunnett test at 0.05 level). **f,** Mechanical stress for each cell type from roots under compacted and non-compacted conditions. Each point corresponds to a cell wall and the horizontal line to the median. Asterisks indicate whether two distributions are significantly different based on a Mann-Whitney test (two-tailed at 0.05 level). Stress inference was performed with ForSys^24^. For the purpose of visualisation, the outliers are not shown. **g-h,** Cortical (g) and epidermal (h) cell wall stiffness of Nipponbare (Nip), *ein2*, and *eil1eil2* under dense media conditions. Two radial sections per genotype were analyzed, with the experiment repeated four times (One-way ANOVA with Dunnett test at 0.05 level). **i,** Expression of *ARF1*, *CESA5*, *6*, and *8* in the elongation zone of Nip and *ein2* seedling roots grown under normal and dense media conditions for 5 days (n=6 per trail, three biological replicates). Statistics presented are from one representative trial (Welch’s t-test with Bonferroni- Dunn method at 0.05 level). **j,** Soil compaction, and associated ethylene accumulation, induces *OsARF1* expression mainly in the cortex cells, leading to suppressed primary wall *CESA* expression. The subsequent reduction in cellulose biosynthesis lead to radial expansion of cortex cells, and thus root swelling, due to reduced cell wall thickness around cortex cells. By contrast, the epidermal wall thickness, and strength, is maintained to support the penetration ability of rice roots in compacted soils. The black line denotes cell wall thickness.

Our results reveal that moderate radial swelling of cortex cells enhanced root growth in compacted soil. This may seem counterintuitive, as cell wall softening and isotropic cell swelling should lead to a lowered directed cell expansion and therefore presumably reduce the ability to penetrate compacted media. However, the epidermal cell layer is also crucial for roots to grow and thrive in soil. For instance, thicker walls of outer small cortical cells in wheat and maize seedling roots enhanced root penetration ability^23^. Indeed, we found that the WT, OE-ARF1 and *cesa6* mutants all contained thick epidermal cell walls when grown on compacted media (Extended Data Fig. 10a,b), with little difference across the genotypes. The epidermal wall stiffness was not significantly different between WT and *arf1-1*, and only slightly lower in OE-ARF1 and *cesa6* (Extended Data Fig. 10c). These observations suggest that a “stiff epidermis- soft cortex” cell wall model facilitates root adaptation to soil compaction. To validate this model, we assessed the mechanical stress in various root cell types from compacted and non-compacted conditions by performing mathematical stress inference using the ForSys^24^ algorithm. The spatial profiles of stresses within root cross sections showed qualitative differences between compacted and non-compacted roots, with cells in the cortex showing less stress in compacted soils than in non-compacted scenarios, while epidermal cells show the opposite trend (Extended Data Fig. 11). Quantitative comparison of the mechanical stresses in the homotypic cell walls revealed that while the sclerenchyma cells were not significantly affected by soil compaction, roots exposed to compacted soils resulted in reduced stresses in the cortex and increased stresses in the epidermal, exodermal and endodermal cells (Fig. 4f). Our results show that under soil compaction, stress accumulates in the outer rims and endodermis and decreases in the cortex.

### Compaction stress triggers *OsARF1* upregulation in an ethylene dependent manner

Our studies reveal that root compaction responses are controlled by OsARF1 repressing *CESA* expression and cell wall thickness in cortical cells, but it is unclear how OsARF1 is regulated by compaction stress. Soil compaction reduces diffusion of the gaseous hormone ethylene, causing this signal to accumulate around plant roots and trigger an ethylene response. This in turn up-regulates levels of auxin and abscisic acid (ABA) signals, which act to repress root elongation and promote root radial expansion, respectively^5,6^. We, therefore, investigated if any of these hormones control *OsARF1* expression. To address this, we grew the *ProOsARF1*:GUS reporter line in control, 10 μM IAA (Auxin), 25 μM ABA or 100 μM ACC (ethylene precursor) conditions. ACC treatment strongly induced *OsARF1* expression, similar to the induction observed in dense agar media, while IAA and ABA showed no or little effects, respectively (Extended Data Fig. 12a). This finding suggests that ethylene may be the primary signal linking compaction and *OsARF1* expression. We also noted moderate induction of *OsARF1* by ABA (25 µM) in the cortical region, though less pronounced than the ACC treatment. Subsequent GUS quantification and RT-qPCR analyses confirmed significant upregulation of *OsARF1* expression under ACC treatment, but showed reduced expression with ABA treatment (Extended Data Fig. 12b,c). This reduction in ABA-treated samples might reflect changes in overall root *OsARF1* expression, as RT-qPCR measurements encompassed whole roots, potentially masking localized expression increases observed in our GUS analyses. These results support ethylene as an inducer of *OsARF1* expression. We hypothesize that ABA may function downstream of ethylene, as suggested by previous studies^5^, partially modulating *OsARF1*, though this relationship requires further investigation.

Given that *OsEIN2*, *OsEIL1*, and *OsEIL2* encode key components of the ethylene response pathway, specifically a response effector and upstream transcription factors^25^, respectively, we analyzed whether *OsARF1* acts downstream of these genes by growing Nip (*Nipponbare*)*, ein2 and eil1eil2* under normal and dense media conditions. Under dense media conditions, *OsARF1* expression was significantly induced in both Nip and *eil1eil2* mutants, but not in *ein2* (Fig. 4i). This suggests that *OsARF1* functions downstream of *OsEIN2*. Furthermore, while dense media treatment led to downregulation of cellulose synthase genes (*OsCESA5*, *6*, *8*) in Nip, this response was abolished in *ein2* mutants (Fig. 4i). Interestingly, in the *eil1eil2* background, the three *OsCESA* genes showed distinct responses: *OsCESA5* maintained its downregulation, *OsCESA6* showed no significant change, and *OsCESA8* was upregulated (Fig. 4i). These differential responses highlight the complex regulatory network connecting ethylene signaling and cellulose synthesis. Interestingly, *ein2* and *eil1* seedling roots, despite their slimmer morphology, successfully penetrate compacted soils^6^, suggesting that ethylene-insensitive roots have enhanced soil penetration capacity. This observation appears to contrast our finding that OE-ARF1 and *cesa6* mutant roots, which are thicker than WT, also show improved soil penetration. To further explore these apparent inconsistencies, we conducted growth experiments where we germinated and grew WT, *ein2* and *eil1eil2* on different media conditions with or without 150 pM Indaziflam. The resulting root thickening (Extended Data Fig. 13) suggests that cellulose synthesis modifications may occur downstream of OsEIN2, OsEIL1 and OsEIL2 to enhance radial swelling in rice roots, though the relationship between these pathways requires further investigation. Notably, *ein2* and *eil1eil2* mutants displayed epidermal and cortical wall stiffness patterns consistent with our model (Fig. 4g,h). We hypothesize that the absence of cell swelling response in *ein2* mutants under compacted conditions may be attributed to ABA accumulation defects^5^. Collectively, these data support that a “stiff epidermis-soft cortex” model promotes root penetration capability in compacted soil conditions.

## Conclusion

Our study provide new conceptual insights into how roots adapt to soil compaction stress through cortical cell expansion, demonstrating how a “stiff epidermis-soft cortex” model effectively responds to soil compaction (Fig. 4f). Under compaction stress and the resulting ethylene response, *OsARF1* expression is induced in the cortex of the elongation zone (Fig. 3a, Extended Data Fig. 12a), which suppresses *OsCESA6* (Extended Data Fig. 6) and other primary wall *CESAs*, in the cortex leading to thinner cell walls and radial expansion of cortex cells (Fig. 4j). The radial expansion of cortex cells exerts pressure on the outer epidermal cell layer, potentially providing axial stability to the root. The increased root diameter, combined with maintained epidermal strength, may enhance stability against ovalization and reduce buckling susceptibility^26^. This “stiff epidermis-soft cortex” model parallels engineering principles used in pipe design, where maintaining perimeter strength while increasing diameter improves structural stability. While OsARF1-mediated cellulose regulation affects root adaptation to compaction, we observed that epidermal cell wall thickening still occurred in WT, OE-ARF1, and *cesa6* under compacted soil conditions (Extended Data Fig. 10a,b). This suggests multiple mechanisms contribute to the ‘thick and stiff epidermal cell wall’ model. Interestingly, OsEIL1 promotes epidermal wall thickening through direct regulation of OsCSLC2-mediated xylan biosynthesis^27^.

In our study, root expansion primarily resulted from cortical cell expansion rather than changes in cell number, which has been identified as a distinct phene in maize roots^28^. Indeed, our analysis of cortical cell numbers in rice under various soil conditions revealed no consistent differences. The relationship between mechanical stress and cellular organization is closely linked to the arrangement of microtubules that coordinates directions of cellulose microfibrils in the cell walls. Indeed, the weakened cortex cell walls may cause an increased pressure on the epidermal cell layers, which in turn could affect, perhaps hyper-align, the microtubules in the epidermis. This could in turn cause an increase in transverse microfibrils that would strengthen the epidermis to prevent further radial swelling of the roots and to aid directed growth of the roots. While challenging in rice root cells, imaging of microtubules in root adaptation to soil compaction remains an important area for future research^29^.

Our research establishes a crucial link between ethylene signaling and root mechanics through the regulation of cell wall synthesis and strength, highlighting the dynamic role of cellulose synthesis in root penetration of soil. ABA is also regulated by ethylene during root compaction, and may function as a downstream signal affecting *OsARF1* levels, as evidenced by its induction of *OsARF1* expression in cortex layers (Extended Data Fig. 12a). In addition, brassinosteroids (BRs) regulate hypocotyl elongation through epidermal wall loosening rather than wall thickness^30^, and BR- responsive genes *HOMEOBOX FROM ARABIDOPSIS THALIANA 7 (HAT7)* and *GT- 2-LIKE 1 (GTL1)* express in cortex to mediate cell elongation by activating cell wall-related genes^31^. However, we did not observe any significant trends in BR gene expression under different soil conditions (Zhu et al., co-submitted)^32^. To sum up, our findings have significant implications for developing crop varieties better adapted to challenging soil conditions through targeted modification of cell wall properties and hormone responses.

## Methods

### Plant material construct development and growth conditions

The background/wild type (WT) of *osarf1*, OE-ARF1, *cesa6* and NG-CESA6 in this study was Rice (*Oryza sativa ssp. japonica*) variety 9522. *Nipponbare* (Nip), *ein2*, and *eil1eil2* were sourced from Rongfeng Huang’s group. The *osarf1*, *cesa6* and *arf1cesa6* knock out alleles were generated using CRISPR/Cas9 technology, with mutant genotyping conducted through PCR amplification of the target regions followed by sequencing of the products. We amplified *OsARF1* coding (2100 bp) and promoter (2915 bp in front of ATG) sequences and then cloned them into pTCK303 and pCAMBIA1301 to produce OE-ARF1 and *ProOsARF1*:GUS, respectively. We also amplified *OsCESA6* promoter (2877 bp upstream of ATG) sequence and then cloned the segment into pCAMBIA1301 to produce *ProOsCESA6*:GUS. mNeonGreen was fused to the *OsCESA6* genomic sequence under the control of the *OsCESA6* promoter and was cloned into pCAMBIA1301 to create NG-CESA6, as described in reference^15^. All plasmids were constructed using the In-Fusion HD Cloning Kit (Takara, Shiga, Japan) and were verified by sequencing at BGI, Beijing, China. These plasmids were introduced into the calli of 9522 or *cesa6* via Agrobacterium (EHA105)-mediated transformation, with hygromycin B used for selection as outlined in reference^32^. All primers used are listed in Extended Data Table 1.

All rice samples were grown and collected at the paddy field of Shanghai Jiao Tong University, under natural conditions from June to October. Seedlings were cultivated in a plant incubator at 28°C with a 16-hour light/8-hour dark cycle and 50%-70% humidity.

### Soil compaction details

Loamy sand from the Newport series (comprising 83.2% sand, 12.1% clay, and 4.7% silt; with 2.93% organic matter and a pH of 6.35; classified as FAO Brown Soil) and clay soil were collected from the University of Nottingham farm at Bunny, Nottinghamshire, UK (52.52°N, 1.07°W). Both soils were passed through a 2 mm sieve. The moisture content was determined by drying the soil at 45°C until a constant weight was achieved.

Mesocosms were prepared by packing them with the sieved soil to achieve a bulk density (BD) of 1.2 g cm^-3^, to simulate a non-compacted condition, and 1.6 g cm^-3^ for a compacted condition. Rice seeds were sterilized with a 25% bleach solution for 10 minutes and subsequently rinsed six times with sterilized water. The seeds were then placed on filter paper for 48 hours in a dark incubator chamber to allow for germination. One germinated seedling per mesocosm was positioned on the soil surface and covered with a 1 cm top layer of soil at 1.2 BD, in both non-compacted and compacted soil conditions. The seedlings were then placed in a rice growth chamber, which was maintained at 28°C, with a 12-hour photoperiod and 70% relative humidity.

### X-Ray CT imaging

Five-day-old seedlings of WT, *osarf1*, and *cesa6* were grown in 3D-printed columns (33 mm in diameter and 100 nm in height) filled with sandy loam soil under non-compacted (1.2 g/cm^3^) and compacted (1.6 g/cm^3^) conditions, respectively. They were then imaged using a GE Phoenix v|tome|x M 240 kV X-ray tomography system at The Hounsfield Facility, University of Nottingham. Three-dimensional image reconstruction was carried out using Datos|REC software (GE Inspection Technologies, Wunstorf, Germany). The roots were segmented from the soil using a polyline tool in VGStudioMAX V2.2 (Volume Graphics GmbH, Germany) to demonstrate the root length phenotype. The scanning protocol involved collecting 3240 projection images in FAST mode (continuous rotation), with the X-ray tube set to an energy of 140 kV and a current of 200 μA. The detector’s exposure time was 131 milliseconds, and the voxel resolution was 57 μm. Each scan had a duration of 7 minutes.

### Agar experimental details

Agar at concentrations of normal (0.3%) and dense (0.6%) (Sigma-Aldrich-7002, Spain) was boiled and poured into tanks measuring 7.5 cm in diameter and 8.5 cm in length to simulate non-compacted and compacted soil conditions, respectively. After the agar solidified, germinated seeds were placed on the media surface, followed by watering (1cm height). The seeds were then grown in a plant incubator (28°C, 16-h light/8-h dark, 70% relative humidity) for five days. Finally, root phenotypes were imaged using a Nikon camera, and the images were analyzed with ImageJ software.

### Imaging of root tip thickness

Root tips, including the root cap, meristematic, elongation, and differentiation zones (approximately 1 cm of the rice root tip), were washed three times with sterilized water. They were then embedded in 5% melted agarose (LabTop Biotechnology, Shanghai, China). Transverse sections, 50 μm thick, were cut using a Leica Vibratome and imaged with a Leica SP5 confocal microscope utilizing the UV laser.

### RNA isolation and RT-qPCR

Root tips, including the root cap, meristematic zone, elongation zone, and differentiation zones (approximately 1.5 cm of the rice root tip), grown in soil or agar media conditions, were sampled in three biological replicates. Total RNA was extracted using Trizol reagent (Invitrogen, Carlsbad, CA, USA) followed by purification with chloroform/isopentyl alcohol. The cDNA was synthesized using the FastQuant RT Kit with gDNase (Tiangen Biotech, Beijing, China). RT-qPCR was conducted using the SYBR Green Premix Pro Taq HS qPCR Kit (Accurate Biotechnology, Hunan, China). The reaction mixture consisted of 7.5 μL SYBR, 2 μL cDNA, 0.3 μL each of forward and reverse primers, and 4.9 μL double distilled water. The Ubiquitin gene served as the reference for assessing gene expression levels. All RT-qPCR primers are listed in Extended Data Table 1.

### Yeast one hybrid screening

Nine promoter fragments – 753 bp for *OsCESA1*, 1973 bp for *OsCESA3*, 1928 bp for *OsCESA5*, 1964 bp for *OsCESA6*, 1751 bp for *OsCESA8*, 1943 bp for *OsCSI1*, 1872 bp for *OsCSLF6*, 2628 bp for *OsCTL*, and 923 bp for *OsSTL*, as well as a 1738 bp fragment for the *OsHARPIN1* promoter – were cloned into the R4L1pDEST_HIS2 plasmid^33^. The cDNA library encompasses 1,143 rice transcription factors (TFs), representing all known TF families. The appropriate 3AT concentration was determined through autoactivation testing. Subsequently, plasmids were transformed into YM4271 yeast strain (Clontech/TAKARA) using TE/LiAc and PEG solution. Yeast plates were incubated at 30°C for 3-7 days, after which images were captured using a camera.

### Dual-Luciferase assay

Rice seedlings were grown for 7 days post-germination in a plant incubator set to 28°C with a 16-hour light/8-hour dark cycle and 50%-70% humidity. Seedling shoots were harvested and digested in an enzyme solution containing 0.2 g Cellulase ‘Onozuka’ RS (Yakult), 0.05 g Macerozyme R10 (Yakult), and β-mercaptoethanol to a final concentration of 10 mM. After degassing twice, the solution was incubated at 28°C and 80 rpm for 2 hours. Protoplasts were isolated using W5 solution (150 mM NaCl, 125 mM CaCl2, 5 mM KCl, 2 mM MES, pH 5.7) and MMg solution (400 mM mannitol, 15 mM MgCl2·6H2O, 4 mM MES, pH 5.7), examined under a microscope, and adjusted to the appropriate concentration (2.0-2.5 x 10^5 cells/ml). The *OsARF1* and *eGFP* coding sequences, serving as effectors, were cloned into the pZmUBQ1p_SX_HSPG plasmid. Similarly, promoter sequences of cellulose synthesis genes, acting as reporters, were cloned into the pGL4.1HSP plasmid. Both effectors and reporters were introduced into the protoplasts using PEG/Ca^2+^ and incubated for 16-18 hours at 28°C with 80 rpm agitation. The relative reporter activity was determined by normalizing the reporter luciferase value to the Renilla luciferase reference value. The incubated protoplasts were lysed using the Dual-LUC reporter assay kit (TOYOINK) and the luminescence was measured using a multi-scan spectrometer (TECAN, Männedorf, Switzerland).

### EMSA

First, we amplified the full-length coding sequence of *OsARF1* and cloned it into the pSP6-T7 vector (Promega, Madison, WI, USA). The OsARF1 protein was then expressed using an in vitro translation kit (TNT T7/SP6 Coupled Wheat Germ Extract System; Promega, Madison, WI, USA) and analyzed by Western blot. Double-stranded M1, M2, and M3 hot probes were labeled with Fluorescein amidite dye at the 5’ end. For competition assays, we used 200-fold excess of non-labeled probes (wt) and labeled mutated probes (mut). The protein-probe binding reaction mixture, which included 250 mM Tris-Acetate, 10 mM DTT, 1 mg/mL BSA, and 20 mM MgAC2, was incubated at 25°C for 20 minutes. The reaction products were then resolved on a 6% native polyacrylamide gel and visualized using the Cy2 channel of a ChemiDoc MP imaging system (BioRad, Hercules, CA, USA).

### ChIP-qPCR

The ChIP assay was conducted in accordance with the procedure described by Zhu et al. (2022), with slight modifications. Approximately 2 g of root tips (about 1.5 cm in length) from WT and *ProOsARF1*:gOsARF1-GFP plants were crosslinked with 1% (v/v) formaldehyde in extraction buffer (0.4 M sucrose, 10 mM Tris-HCl, 5 mM β- mercaptoethanol, 0.1 mM PMSF, and a protease inhibitor cocktail, pH 8.0), then pulverized in liquid nitrogen. Chromatin was subsequently isolated and sonicated to generate DNA fragments ranging from 200 to 500 bp. GFP-Trap Magnetic Agarose (ChromoTek, Munich, Germany) was utilized to precipitate OsARF1-DNA complexes. Fragments of the *OsCESA5* and *OsCESA6* promoters were quantified by RT-qPCR using primers listed in Supplemental Table S1. Enrichment in the *ProOsARF1*:gOsARF1-GFP samples was compared to the levels in WT plants.

### Hormone and Indaziflam treatment conditions

Rice seeds of *ProOsARF1*:GUS were sterilized with 70% (v/v) ethanol for 1 minute and 3% (v/v) sodium hypochlorite solution for 15 minutes, followed by at least six rinses with sterile double distilled water, for 3 minutes each. The seeds were then placed on filter paper in germination boxes for 3 days under a 16-hour light/8-hour dark cycle. After germination, the seeds were exposed to various conditions: mock (water), 1 μM IAA, 25 μM ABA, and 100 μM ACC for 2 days.

Similarly, germinated seeds of WT, *osarf1* and OE-ARF1 were transferred to black boxes containing growth medium supplemented with either 150 pM DMSO or 150 pM indaziflam, and 250 pM DMSO or 250 pM indaziflam, and 50 nM DMSO or 50 nM DCB. These were incubated in a plant incubator at 28°C, with a 16-hour light/8-hour dark cycle and 50%-70% humidity for five days. The primary root lengths of the mock-treated and experimental groups were measured and compared using ImageJ. This experiment was performed in triplicate.

### GUS histochemical and quantification

Seedlings of the rice auxin response factor transcriptional reporter (*ProOsARF1*:GUS and *ProOsCESA6*:GUS) were grown in conditions including mock (water with added DMSO), 0.6% agar media, 1 μM IAA, 25 μM ABA, and 100 μM ACC. Root tips were then collected and incubated overnight in GUS buffer (50 mM NaPO4 buffer, pH 7.0, 10 mg/mL X-Gluc, and 0.02% (v/v) Triton X-100) at 37°C. Subsequently, the samples were washed with 70% (v/v) ethanol until they became transparent. Cross-sections were prepared using a Leica vibratome at a thickness of 50 μm. Images were taken with a Leica light microscope (M205A) equipped with a CCD camera, and additional photographs were obtained using a Nikon H600L light microscope (Tokyo, Japan).

For GUS activity quantification, approximately 0.1 g of *ProOsARF1*:GUS roots (1.5 cm) were harvested after 2-day treatment with mock (DMSO), 1 μM IAA, 25 μM ABA, or 100 μM ACC. Root tissues were ground in liquid nitrogen and homogenized in 1 mL GUS extraction buffer (1 M Na2HPO4, 1 M NaH2PO4, 10% SDS, 0.5 M EDTA, Triton X-100, β-Mercaptoethanol). The homogenate was centrifuged at 12,000 rpm for 5 min at 4°C, and protein concentration in the supernatant was determined using the Bradford method. For the GUS activity assay, 100 μL of protein extract was mixed with 400 μL pre-warmed (37°C) GUS extraction buffer and 500 μL MUG substrate (2 mM). The reaction mixture was incubated at 37°C, and 200 μL aliquots were collected at 0, 15, 30, 45, and 60 mins intervals. Each aliquot was immediately mixed with 800 μL stop solution (0.2 M Na2CO3) and stored at room temperature in the dark. Fluorescence intensity was measured using a fluorescence microplate reader (excitation: 365 nm, emission: 455 nm, slit width: 10 nm). The rate of fluorescence intensity change was calculated from the slope of the fluorescence intensity versus time plot and normalized to total protein content to determine specific GUS activity.

### Cellulose measurements

Cellulose content was quantified following Kumar and Turner (2015) with modifications. Five-day-old seedlings of wild-type, *osarf1*, and OE-ARF1 were grown in 0.3% or 0.6% agar media (Sigma-Aldrich-7002, Spain) at 28°C under a 16/8-hour light/dark cycle. Root tips (approximately 1.5 cm) were harvested and processed to obtain alcohol-insoluble residue (AIR). Briefly, samples were incubated in 1.5 mL 70% ethanol at 70°C for 1 hour, followed by a second 45-minute incubation with fresh ethanol. After ethanol removal, samples were treated with 1 mL acetone at room temperature for 2 minutes. The resulting AIR was dried, weighed, and transferred to 15 mL Falcon tubes. For cellulose extraction, 3 mL of acetic/nitric reagent was added to each sample (including a blank control), and tubes were heated in a boiling water bath for 30 minutes. After cooling, samples were centrifuged at 2,000 × g for 10 minutes. The supernatant was carefully discarded, and pellets were washed with 8 mL water, gently resuspended, and incubated for 15 minutes before centrifugation (2,000 × g, 10 minutes). A final wash was performed with 4 mL acetone, followed by a 5-minute incubation and centrifugation (2,000 × g, 10 minutes). Samples were dried in a vacuum oven at 30-40°C. The dried samples were hydrolyzed in 1 mL of 67% sulfuric acid with shaking (180 rpm) at room temperature until complete dissolution. For colorimetric determination, 40 μL of each hydrolysate was mixed with 1 mL ice-cold anthrone reagent and heated in boiling water for 5 minutes, followed by immediate ice quenching. Cellulose content was calculated as percentage of cell wall (CW) per AIR using glucose standard curves according to the formula: % CW = (μg glucose)/(μg AIR) × 100 × (H₂SO₄ volume)/(sample volume).

### Cell wall thickness experimental details

Roots from 5-day-old seedlings of WT, *osarf1*, OE-ARF1, and *cesa6* were cultivated in 0.3% and 0.6% agar media (Sigma-Aldrich-7002, Spain) at 28°C under a 16/8-hour light/dark cycle. Root tips, approximately 1 cm in length, were harvested and fixed in a solution of 3% (w/v) paraformaldehyde and 0.25% glutaraldehyde in 0.2 N sodium phosphate buffer (pH 7.0). These root tips were then post-fixed in 2% OsO4 in PBS (pH 7.2). The samples were dehydrated through a graded ethanol series (70% for 30 minutes, 90% for 30 minutes, and 100% three times, each for 30 minutes) and then transitioned through ethanol/epoxypropane mixtures (2:1, 1:1, and 1:2), followed by pure epoxypropane, each for 10 minutes. Samples were embedded in acrylic resin (London Resin Company) and placed in a drying oven at 37°C for 5-12 hours to eliminate bubbles, then the temperature was increased to 45°C for 2 hours, and finally to 65°C for 48 hours. Ultra-thin sections (50 to 70 nm) were double-stained with 2% (w/v) uranyl acetate and 2.6% (w/v) lead citrate aqueous solution and examined with a JEM-1230 transmission electron microscope (JEOL) at 80 kV.

### Cell wall stiffness experimental details

Roots from 5-day-old seedlings of WT, *osarf1*, OE-ARF1, and *cesa6* were grown in 0.6% agar media (Sigma-Aldrich-7002, Spain) at 28°C with a 16/8-hour light/dark cycle. Primary root tips, measured to 1.5 cm, were embedded in 5% melted agarose (LabTop Biotechnology, Shanghai, China) for sectioning. Longitudinal sections of 50 µm thickness were produced using a Leica vibratome (frequency 50 Hz, amplitude 1 mm), and light microscopy was employed to ensure the stele and cortical tissues were correctly positioned with the elongation zone visible. The sections were then stored in deionized water at 4°C overnight.

AFM assays were conducted following the methods described by Fusi et al. (2022). A Dimension ICON (Bruker Nano) equipped with NanoScope Analysis 9.4 software was utilized to probe all root samples. The MLCT-C probe (Bruker Nano), with an average spring constant of 0.01 N/m, indentation depth (varying from samples, 200-300 nm), speed (3000 nm/s), and force limit (300 pN) was used in the experiments. Root sections set in agarose were affixed to glass slides with tape and hydrated with deionized water for 30 minutes prior to AFM analysis. While operating in force-spectroscopy mode under water-hydrated conditions, ten to fifteen independent areas within the visible cortex and epidermis zones (as shown in Fig. 4d) were examined. Four biological replicates were conducted for each plant line. Apparent stiffness (pN/nm) values were extracted from individual force-distance curves using a contact point based fit and a linear stiffness model, using NanoScope Analysis 1.8 software.

### Image segmentation and cell type classification

We imaged root cross-sections (100 µm thick, position 2-2.5 cm behind the root tip) from compacted and non-compacted soil conditions. The cell walls were stained with Calcofluor White 0.1% aqueous solution for 5 min and washed with DI water. The cross-sections were imaged in a Leica TCS SP5 confocal microscope. We segmented the images using Epyseg^38^ and Tissue Analyzer^39^ to manually curate the images where necessary.

We developed a semi-automated computational pipeline to identify the cells and assign them to the appropriate cell type. We classified the cells into five groups: epidermal, exodermal, sclerenchymatous, cortical, and endodermal. Cells that did not fall into any of these categories or were clearly broken were excluded from analysis together with gas-filled spaces. We created masks to surround the main spatial domains of each cell type. To identify cell types, we calculated the geometric centroid of each cell and created a k-dimensional tree to partition the 2D space using the masks. This allowed us to efficiently find the cells’ centres within a particular type’s mask. Finally, we manually confirmed and corrected the resulting cell type assignments when necessary.

### Mechanical stress inference

To infer the mechanical state of the cell walls, we used ForSys, an algorithm capable of inferring mechanical stress in tissues^24^. ForSys inputs the segmented image and constructs a matrix representation of the root cross section by looking at the vertices where three cell walls meet, which we call pivot vertices. The force assigned to each cell wall is assumed in the direction of a circular fit of the wall at the pivot vertex *i* (*r̂*_*ij*_), with an unknown magnitude (*λ*_*ij*_) representing the stress of the cell wall that joins vertices *i* and *j*:

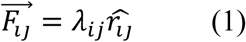

Note that, even though the stress is symmetrical, *i.e.*, *λ*_*ij*_ = *λ*_*ji*_, the direction of the force for a cell wall joining two pivot vertices, *i* and *j,* will differ depending on whether we are performing the calculation from the pivot *i* (*r̂*_*ij*_) or pivot *j* (*r̂*_*ji*_). A force balance is applied at each pivot vertices, giving for pivot *i*

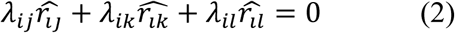

where *j*, *k*, and *l* are the pivots connecting to vertex *i*.

The resulting system of equations has two equations per pivot vertex and one unknown per cell wall. Furthermore, to avoid a trivial solution where all the stresses *λ* are identically null, we added a Lagrange multiplier in which the average value of the stresses should equal one over the entire microscopy image of the root cross-section.

The complete system is further encoded in a geometrical matrix [*M*_*λ*_] as

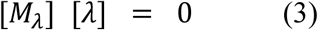

We used the non-negative least squares solver provided in the ForSys tool. The solutions obtained by ForSys are a relative measure of the magnitude of the stress components at each cell wall.

## Acknowledgements

We thank Rongfeng Huang for providing rice *ein2* and *eil1eil2* lines, Ms. F. Tobe, Ms. Kijima Sakutaro and Ms. Kwimi Chuang for their technical contribution to Y1H screens.

## Author contributions

Conceptualization: J.Z., D.Z., S.P.

Methodology: J.Z., Z.Q., J.B.L., Z.Y.L., E.F., L.P.O., A.B., S.S., N.M., X.B.Z, M.Y.Z, J.S.

Investigation: J.Z., B.K.P., N.M., W.L., M.J.B., D.Z., S.P.

Visualization: J.Z., M.J.B., D.Z., S.P.

Funding acquisition: J.Z., B.K.P., D.Z., S.P.

Project administration: M.J.B., D.Z., S.P.

Supervision: B.K.P., D.Z., S.P.

Writing – original draft: J.Z., D.Z., S.P.

Writing – review & editing: J.Z., O.C., N. M., B.K.P., W.L., M.J.B., D.Z., S.P.

## Data and materials availability

No restrictions are placed on materials. Details of all data and materials used in the analysis are available in the main text or the supplementary materials.

## List of Extended Data

Extended Data 1: Yeast-one-hybrid screening results

## Extended Data Figures

**Extended Data Fig. 1:**
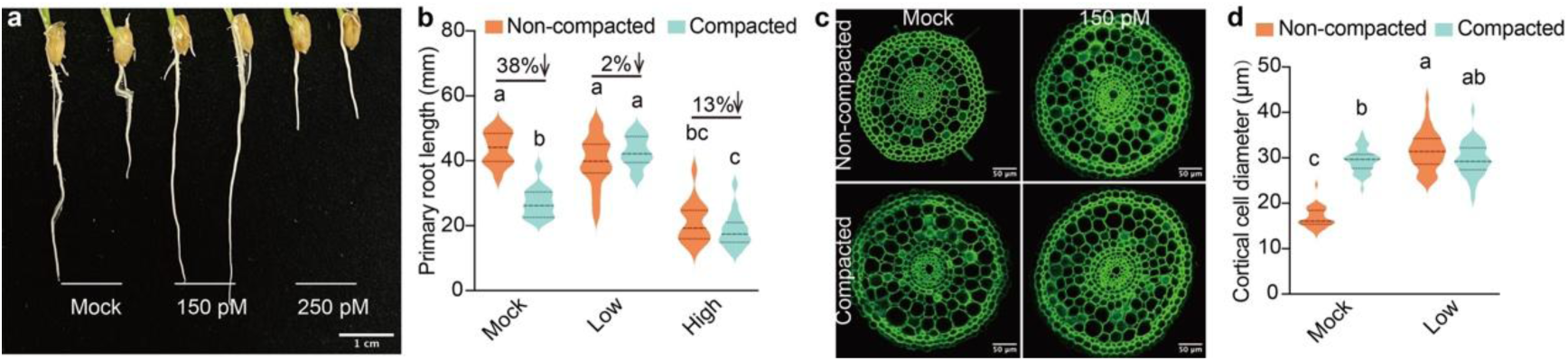
Low concentrations of Indaziflam enhance rice root penetration. **a-b**, Primary root growth of 7-day-old WT seedlings cultured on media supplemented with 150 pM or 250 pM Indaziflam under normal (left panel) and dense (right panel) conditions, with DMSO as the mock treatment. The experiment was conducted in triplicate, yielding consistent trends (n=10 per trial). Statistical analysis was performed using two-way ANOVA and Tukey test, different letters denote significant differences at 0.05 level, *p* (interaction) < 0.0001). Data shown are from one representative trial. Percentages indicate the difference in root length between dense and normal conditions, relative to root length on normal media conditions. Arrows highlight growth reduction in dense versus normal conditions. Scale bar in (a): 1 cm. **c,** Cross-sections of 5-day-old WT roots grown on media supplemented with DMSO (mock) or 150 pM Indaziflam, under normal or dense conditions. Sections (50 µm thickness) were prepared using a vibratome and visualized under UV light (405 nm) with an SP5 confocal microscope. Scale bar: 50 µm. **d,** Cortical cell diameter measurements from sections similar to those in (c). Data represent 40 cells from 5 sections per condition. The experiment was repeated three times with consistent results. Statistical analysis was performed using two-way ANOVA and Tukey test, different letters denote significant differences at 0.05 level, *p* (interaction) < 0.0001).

**Extended Data Fig. 2:**
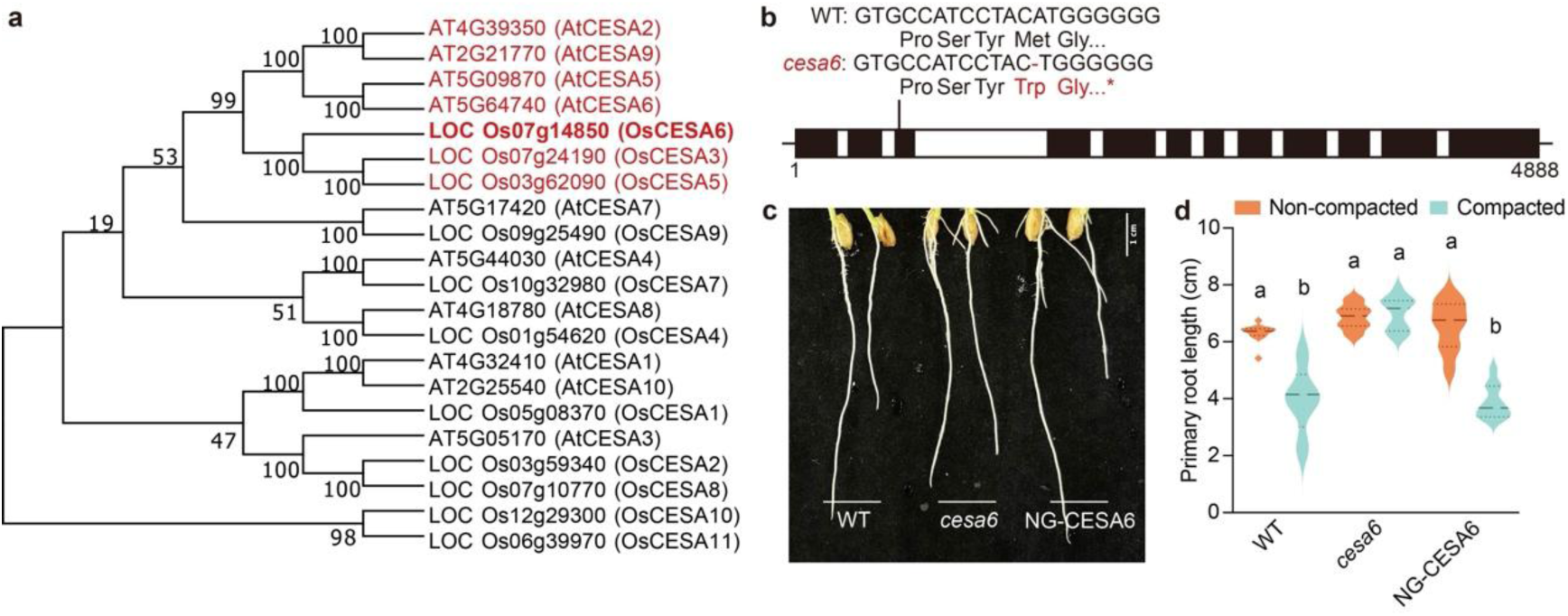
Root penetration phenotype of *cesa6* mutants in agar media. **a,** A phylogenetic tree of the 10 CESA proteins from Arabidopsis and the 11 CESA proteins from rice. AtCESA6 is functionally redundant to CESA2, 5 and 9^40^ implying a similar relationship of OsCESA6 and OsCESA3 and 5. Analysis was performed using MEGA 7.0. Bootstrap values indicate the homology of amino acid sequences. **b,** Genotypic characterization of CRISPR-Cas9 *cesa6* mutant lines revealed a deletion of an ‘A’ in the third exon, resulting in a frameshift mutation and premature termination. Black and white boxes indicate exons and introns, respectively. **c-d,** Primary root penetration phenotypes of WT, *cesa6* mutants, and NG-CESA6 (complementation line of *cesa6*) seedlings grown on normal (left panel) and dense (right panel) agar media for 5 days. The experiment was conducted in triplicate, yielding consistent trends (n=15 per trial). Statistical analysis was performed using two-way ANOVA and Tukey test, different letters denote significant differences at 0.05 level, *p* (interaction) < 0.0001).

**Extended Data Fig. 3:**
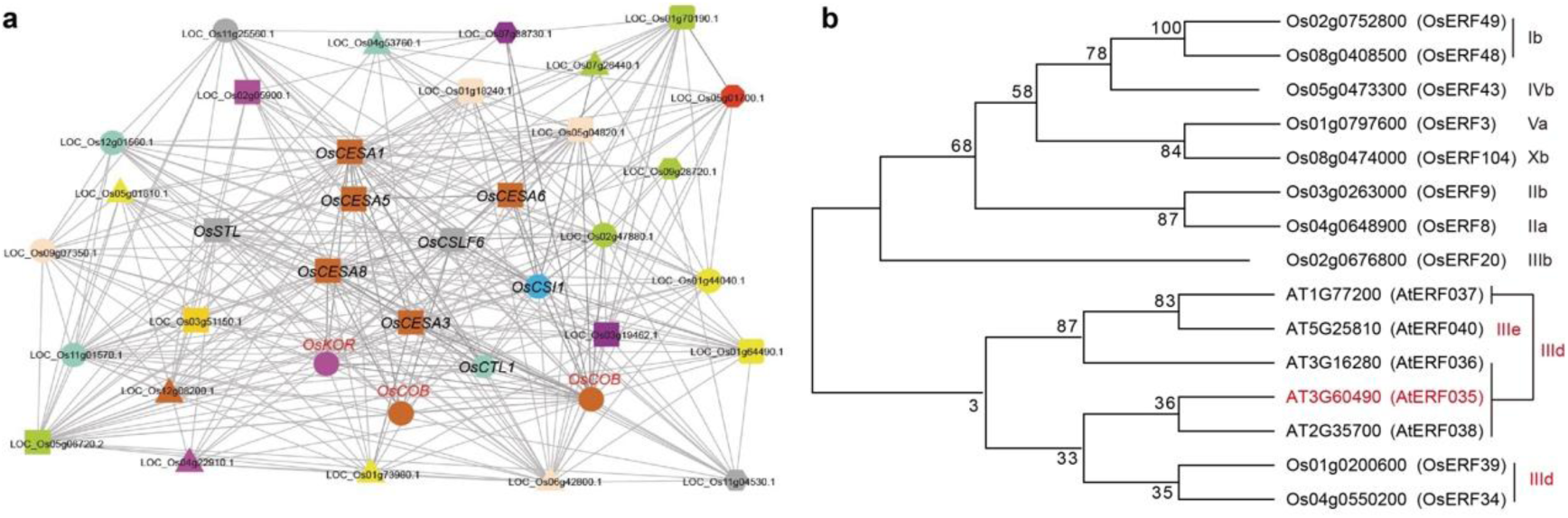
Co-expression network of rice *CESA* genes and ERF transcription factor phylogeny. **a**, Co-expression network centered on primary wall *CESA* genes (represented by orange squares) in rice. Nodes of various colors and shapes correspond to genes from different gene families. Edges indicate significant co-expression relationships between *CESAs* and the genes represented by the nodes. Network is taken from (Mutwil et al., 2011) and truncated. **b**, A phylogenetic tree of selected subclades of rice and Arabidopsis *ERFs*. *OsERF34*, which binds to the promoters of seven cellulose biosynthesis genes, and *OsERF39*, are both close homologs of *AtERF35*, a major primary cell wall regulator in Arabidopsis.

**Extended Data Fig. 4:**
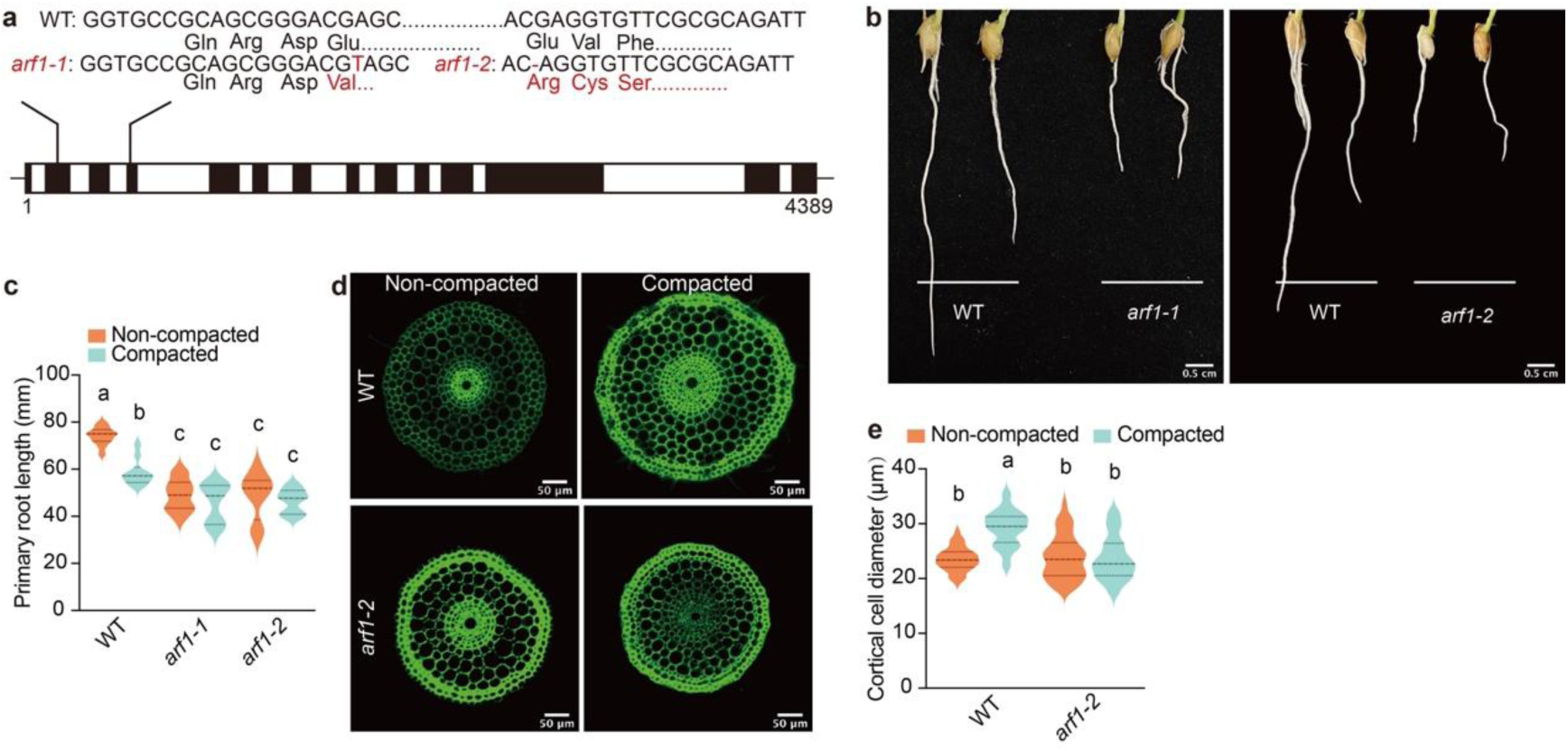
Mutations of OsARF1 cause repression of rice root elongation and swelling in dense media conditions. **a,** Schematic representation of two *arf1* mutant lines: *arf1-1* and *arf1-2*. The *arf1-1* line features a ‘T’ insertion in the second exon, while *arf1-2* has a ‘G’ deletion in the fourth exon. Both mutations induce frameshifts leading to premature termination. Exons and introns are depicted by black and white boxes, respectively. **b-c,** Root penetration phenotypes of WT, *arf1-1*, and *arf1-2* seedlings grown on normal and dense media for 5 days. The experiment was conducted in triplicate, yielding consistent trends (n=12 per trial). Statistical analysis was performed using two-way ANOVA and Tukey test, different letters denote significant differences at 0.05 level, *p* (interaction) = 0.004). **d,** Cross-sections of 5-day-old WT and *arf1-2* (elongation zone) grown under normal and dense media conditions. Sections (50 µm thickness) were prepared using a vibratome and visualized under UV light (405 nm) with an SP5 confocal microscope. Scale bar: 50 µm. **e,** Cortical cell diameter measurements from sections similar to those in (d). Data represent 40 cells from 5 sections per condition. The experiment was repeated three times with consistent results. Statistical analysis was performed using two-way ANOVA and Tukey test, different letters denote significant differences at 0.05 level, *p* (interaction) < 0.0001).

**Extended Data Fig. 5:**
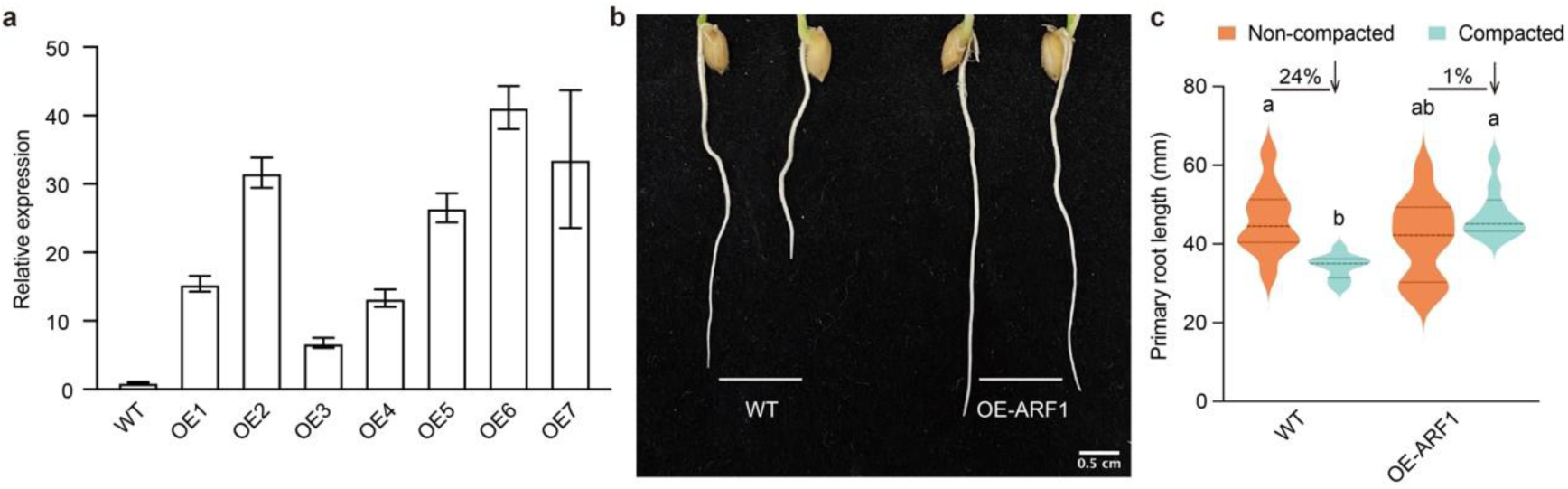
Overexpression of *OsARF1* enhances rice root penetration in dense media. **a,** Expression level of *OsARF1* in seven overexpression lines was quantified using RT-qPCR. Seedling roots were sampled for analysis. The maize ubiquitin gene served as the reference for assessing gene expression levels. **b-c,** Root penetration phenotype and length of WT and OE-ARF1 seedlings grown on normal and dense media conditions for 5 days. The experiment was conducted in triplicate, yielding consistent trends (n=10 per trial). Data shown are from one representative trial. Statistical analysis was performed using two-way ANOVA and Tukey test, different letters denote significant differences at 0.05 level, *p* (interaction) = 0.0008). Percentages indicate the difference in root length between dense and normal conditions, relative to root length in normal conditions. Arrows highlight growth reduction in dense versus normal conditions.

**Extended Data Fig. 6:**
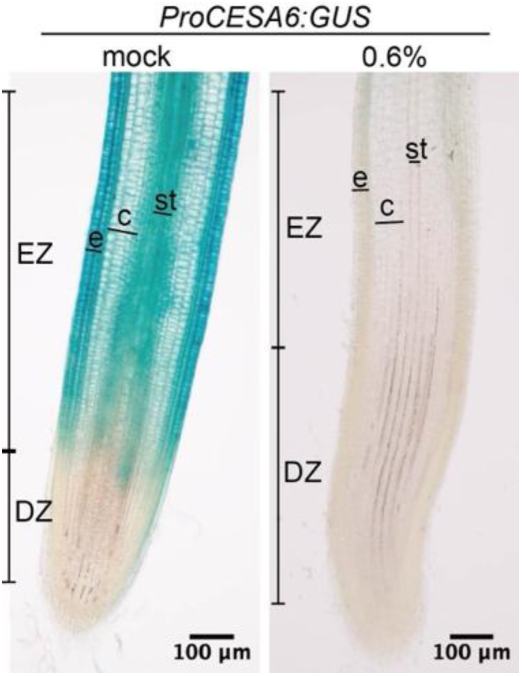
Compacted media reduced *OsCESA6* level in the elongation zone of rice roots. *ProOsCESA6:GUS* lines were cultured in mock (water; left panel) and dense media (right panel) for 48 hours, followed by GUS staining and stereoscopic imaging. Longitudinal sections (50 µm thickness) were prepared using a vibratome. Scale bars: 100 µm. EZ: elongation zone, DZ: division zone, e: epidermis, c: cortex, st: stele.

**Extended Data Fig. 7:**
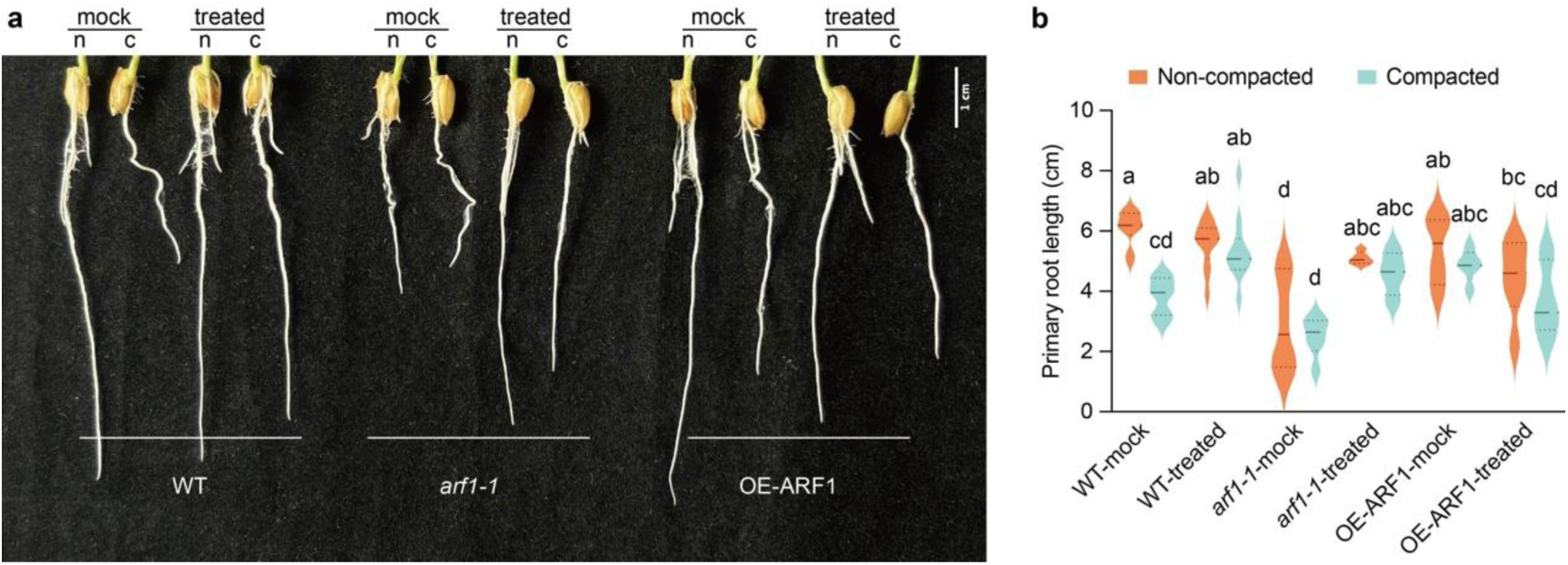
Indaziflam treatment mitigates the root phenotype of *arf1- 1* mutants. **a,** Comparison of primary root lengths in WT, *arf1-1*, and OE-ARF1 seedlings grown for 5 days on normal (n) and dense (c) media. Each group shows roots grown with mock (DMSO, left two roots) and treated (150 pM Indaziflam, right two roots). Scale bar: 1 cm. **b,** Quantification of root lengths from (a). The experiment was replicated thrice, consistently yielding similar trends (n=11 per trial). Statistical analysis was performed using two-way ANOVA and Tukey test, different letters denote significant differences at 0.05 level, *p* (interaction) = 0.0184).

**Extended Data Fig. 8:**
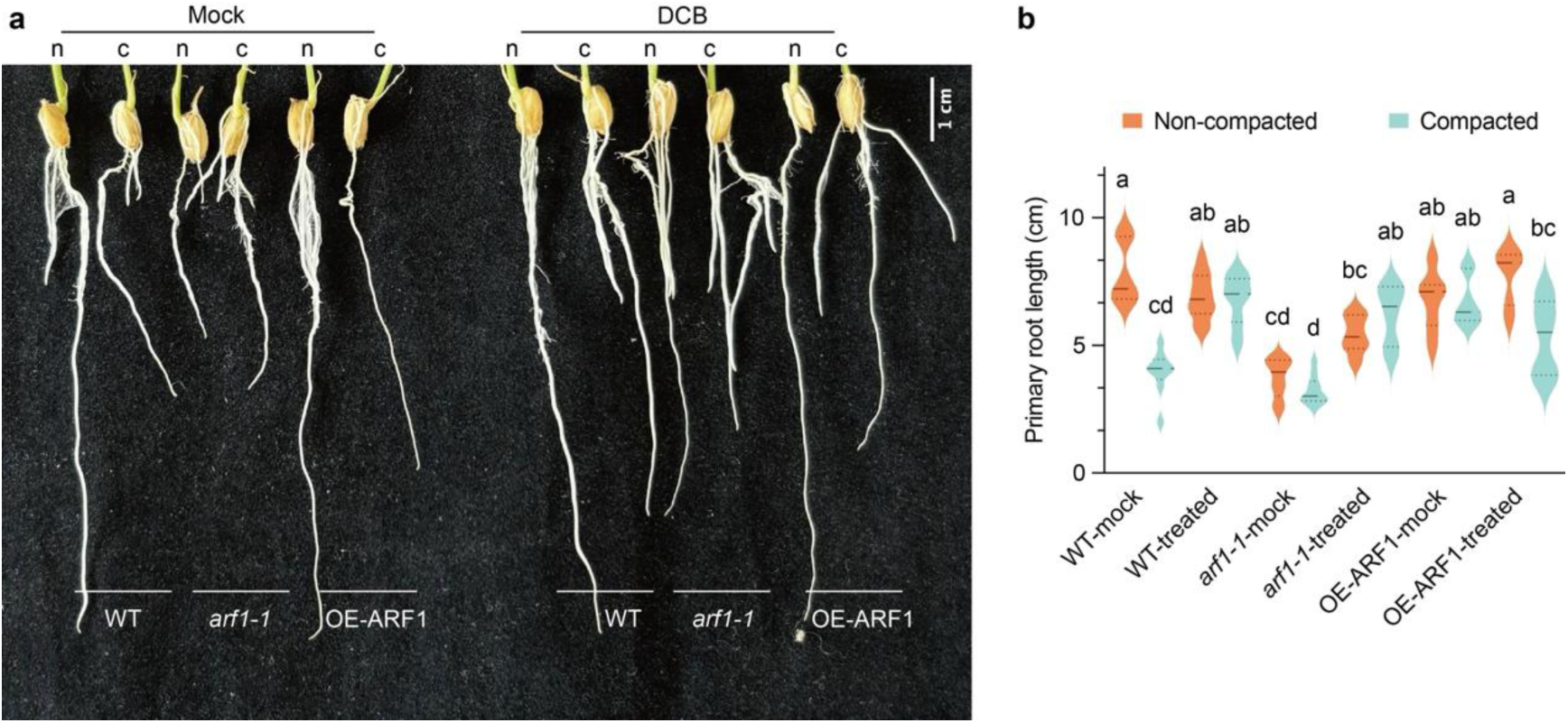
DCB treatment mitigates the root phenotype of *arf1-1* mutants. **a,** Comparison of primary root lengths in WT, *arf1-1*, and OE-ARF1 seedlings grown for 5 days on normal (n) and dense (c) media. Each group shows roots grown with mock (DMSO) and 50 nM DCB treatments. Scale bar: 1 cm. **b,** Quantification of root lengths from (a). The experiment was replicated thrice, consistently yielding similar trends (n=12 per trial). Statistical analysis was performed using two-way ANOVA and Tukey test, different letters denote significant differences at 0.05 level, *p* (interaction) < 0.0001).

**Extended Data Fig. 9:**
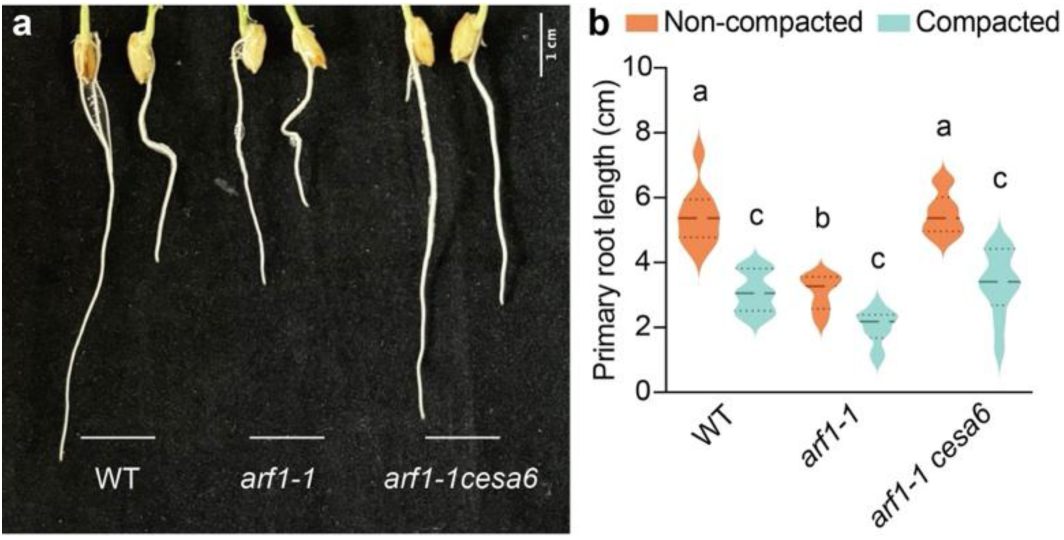
The *arf1* root phenotype is rescued in *arf1cesa6* double mutant line. Root penetration phenotype and length of WT, *arf1-1*, and *arf1cesa6* seedlings grown on normal and dense media conditions for 5 days. The experiment was conducted in triplicate, yielding consistent trends (n=10 per trial). Statistical analysis was performed using two-way ANOVA and Tukey test, different letters denote significant differences at 0.05 level, *p* (interaction) = 0.0156).

**Extended Data Fig. 10:**
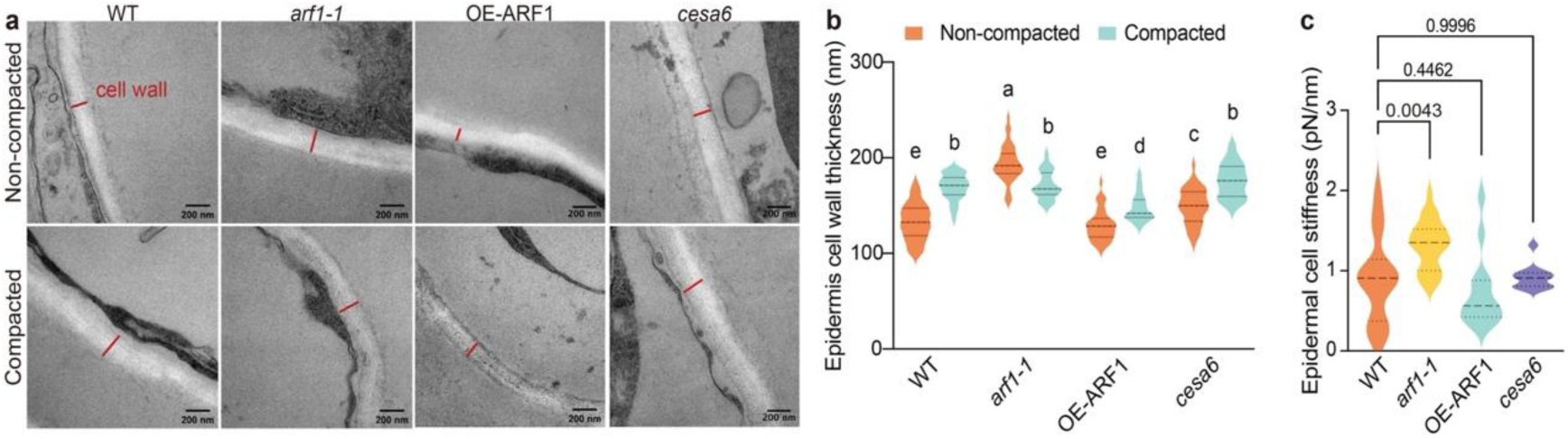
Thickened and stiff epidermal cell walls facilitate root adaptation to soil compaction. **a-b,** Transmission electron microscopy (TEM) images showing cell wall thickness in epidermal cells of the elongation zone from 5-day-old WT, *arf1-1*, OE-ARF1, and *cesa6* seedlings grown on normal or dense media. Measurements were taken from 40 cells across 2 sections per genotype, with the experiment repeated twice. Statistical analysis was performed using two-way ANOVA and Tukey test, different letters denote significant differences at 0.05 level, *p* (interaction) < 0.0001). **c,** Cell wall stiffness in epidermal cells of WT, *arf1-1*, OE-ARF1, and *cesa6* lines grown under dense media conditions. Samples were prepared as 50 µm longitudinal sections using a vibratome. Two sections per genotype were analyzed, with the experiment repeated four times. Statistical analysis was performed using one-way ANOVA with Dunnett test at 0.05 level.

**Extended Data Fig. 11:**
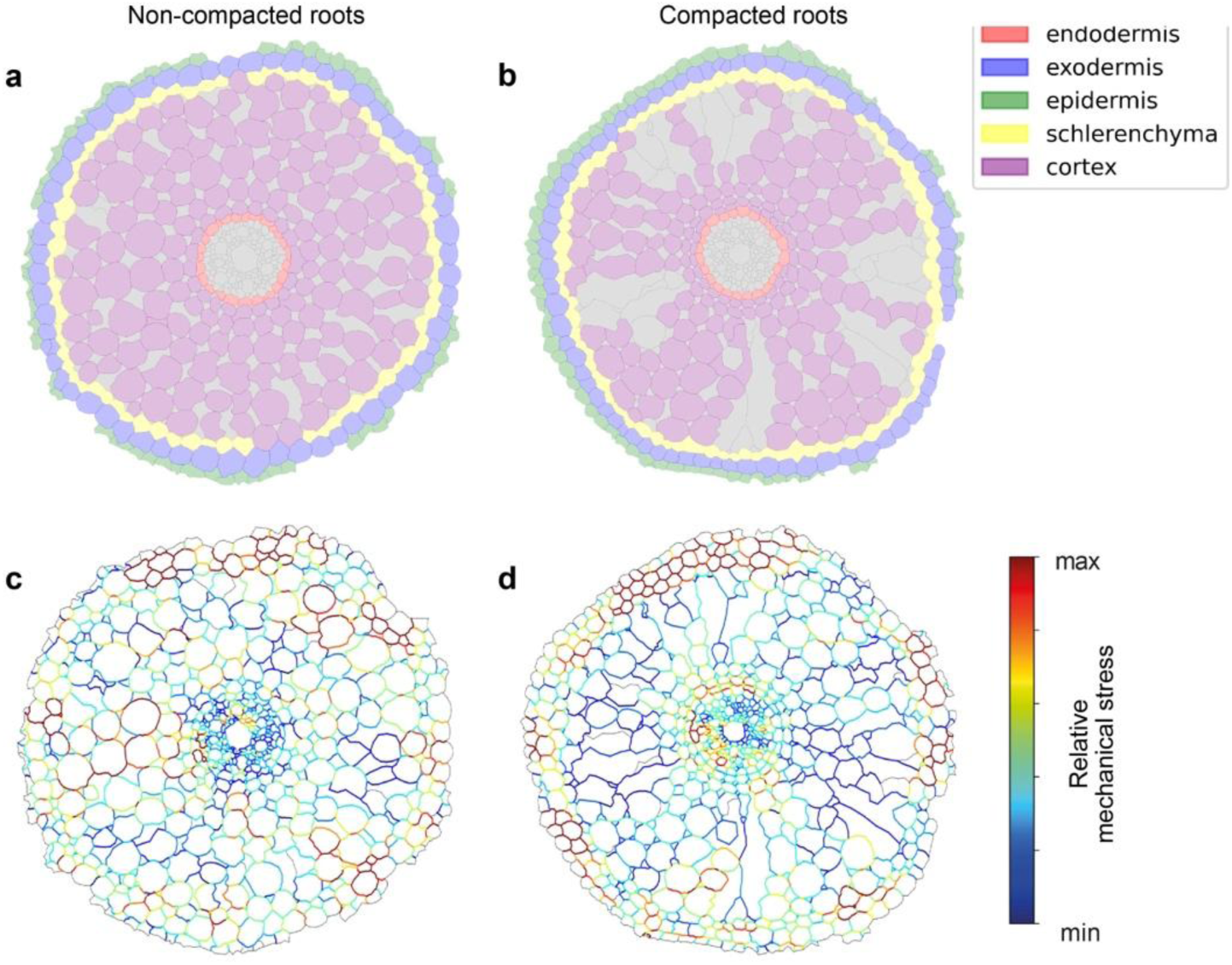
Effect of soil compaction in the mechanical state of the cell walls. **a-b**, Cell types in the root are identified and categorised into five groups, indicated by the cell color. A grey color indicates that the cell did not fall into any of the prescribed groups. The cell contours were obtained through an image analysis pipeline involving image segmentation of the microscopic image of the root cross sections and manual curation. Here, we present two representative examples of a root developed in non-compacted (a) and compacted soil (b). **c-d**, We used the stress inference algorithm ForSys1 to estimate the relative mechanical stress of the tissue, color-coded in the cell walls.

**Extended Data Fig. 12:**
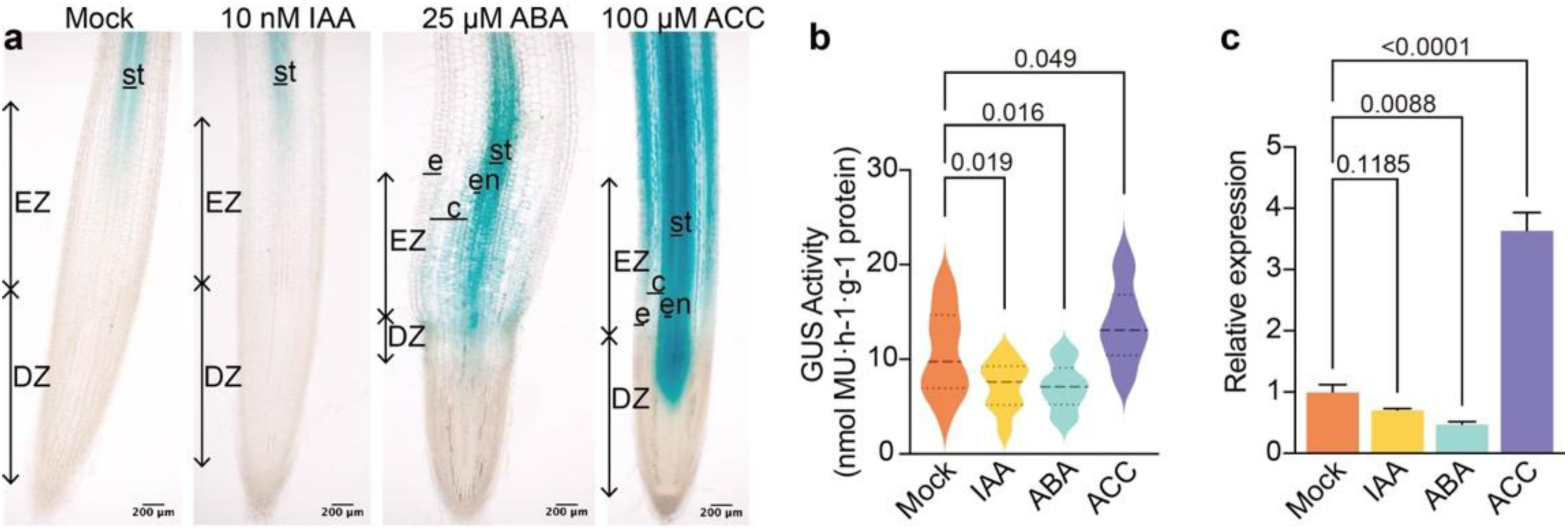
OsARF1 functions downstream of ethylene in soil compaction responses of roots. **a,** *ProARF1*:GUS seedling roots were grown for 48 hours on media supplemented with mock (DMSO), IAA (10 µM), ABA (25 µM), or ACC (100 µM), then stained for GUS activity and imaged using a stereoscope. Longitudinal sections (50 µm thickness) were prepared using a vibratome. Scale bar: 200 µm. EZ: elongation zone, DZ: division zone, e: epidermis, c: cortex, en: endodermis, st: stele. **b,** GUS quantification (change in MU protein amount per unit time) for samples in (a). Each treatment had 16 replicates. Statistical analysis was performed using one-way ANOVA with Dunnett test at 0.05 level**. c,** *OsARF1* expression levels in the elongation zone of WT seedling roots after 48-hour treatment with mock (DMSO), IAA (10 µM), ABA (25 µM), or ACC (100 µM). Samples were collected from six seedling root tips per group, with the experiment repeated three times. Statistical analysis was performed using one-way ANOVA with Dunnett test at 0.05 level. Bars indicate mean ± SD.

**Extended Data Fig. 13:**
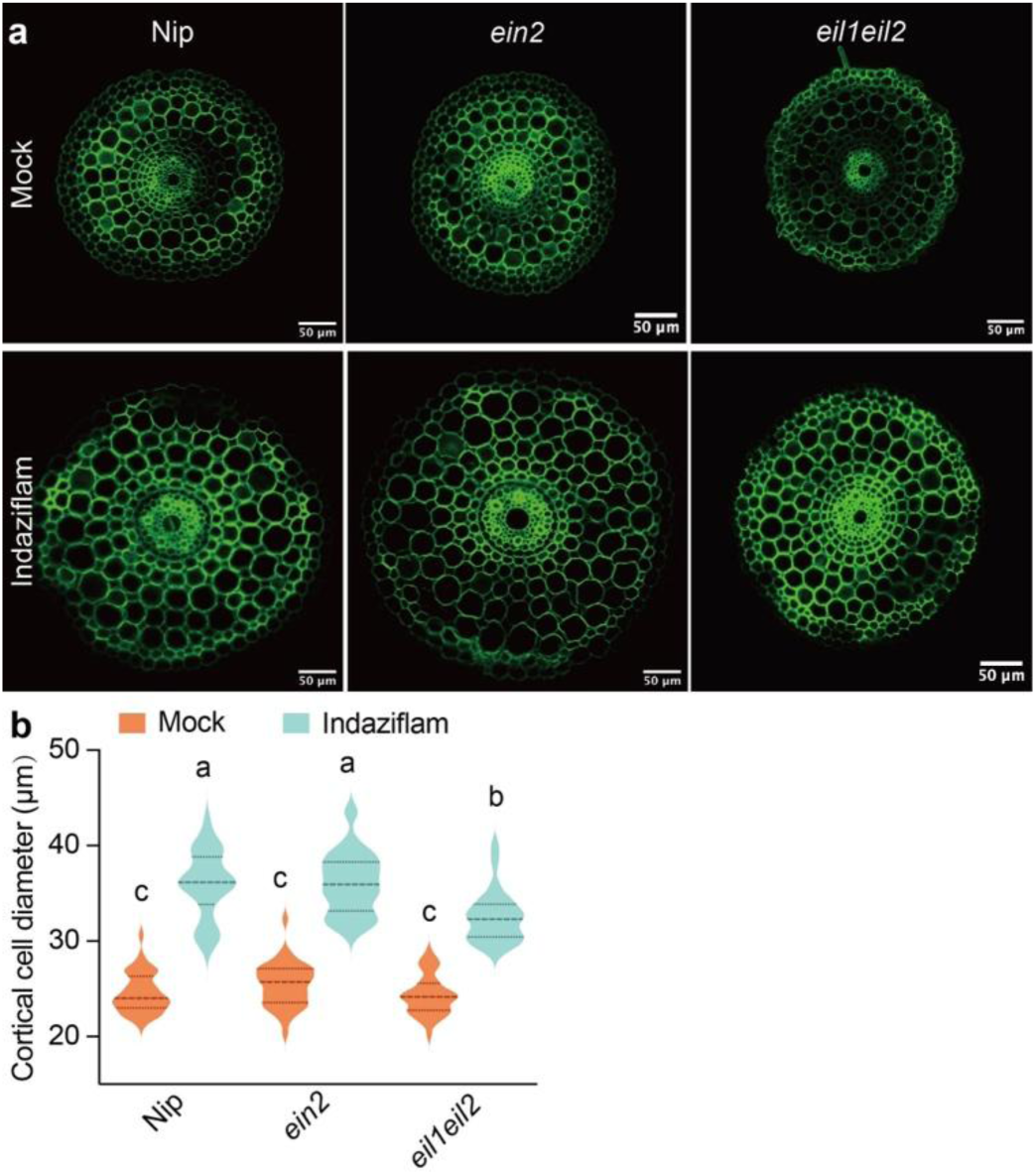
Indaziflam rescues root expansion of *ein2* and *eil1eil2* in compacted conditions. **a-b,** Root sections from the elongation zones of 5-day-old WT, *ein2*, and *eil1eil2* seedlings grown on media supplemented with DMSO (mock) or Indaziflam. Measurements were taken from 40 cells across 5 sections per genotype. The experiment was conducted in triplicate with consistent trends. Statistical analysis was performed using two-way ANOVA and Tukey test, different letters denote significant differences at 0.05 level, *p* (interaction) = 0.0005).

## Extended Data Table

**Extended Data Table 1:**
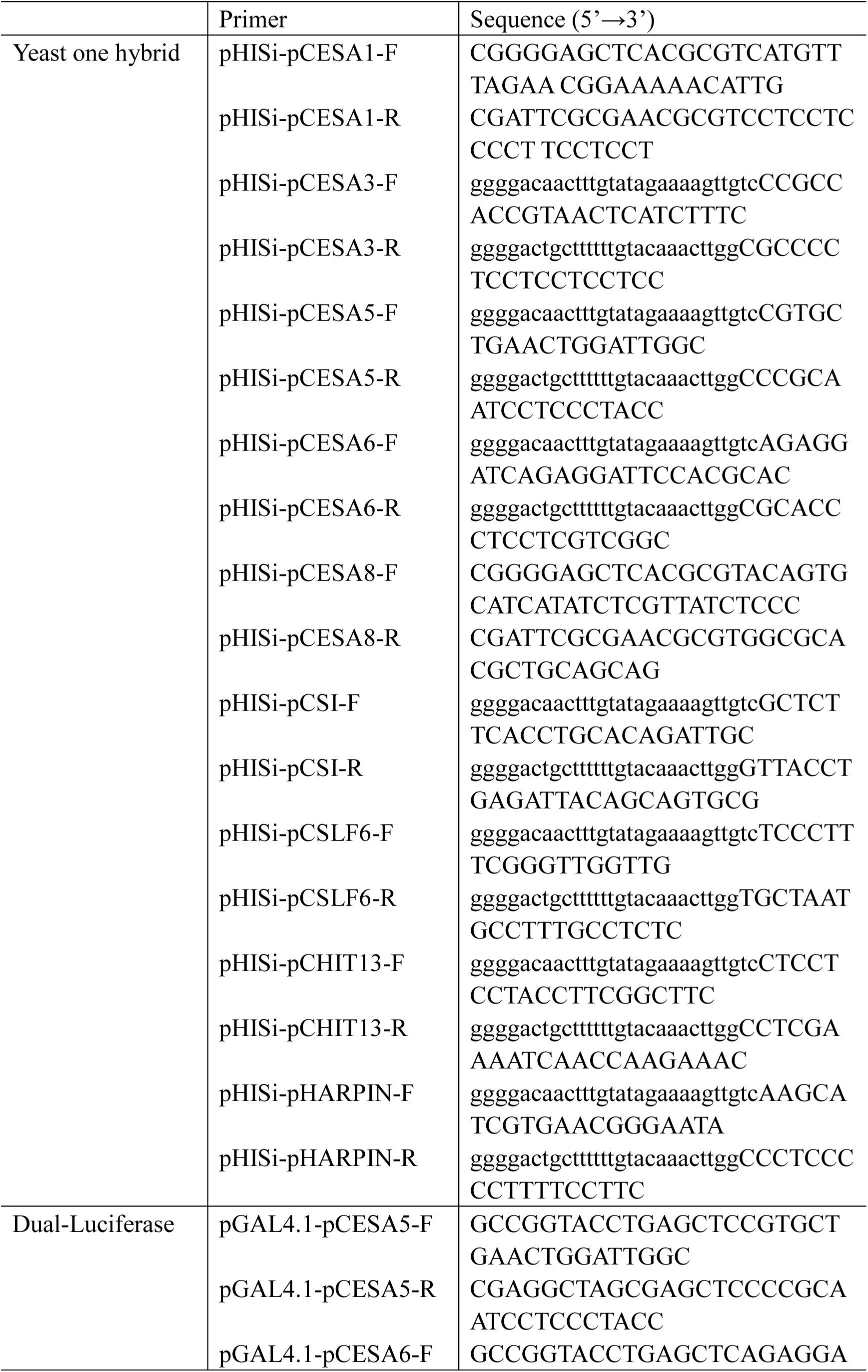
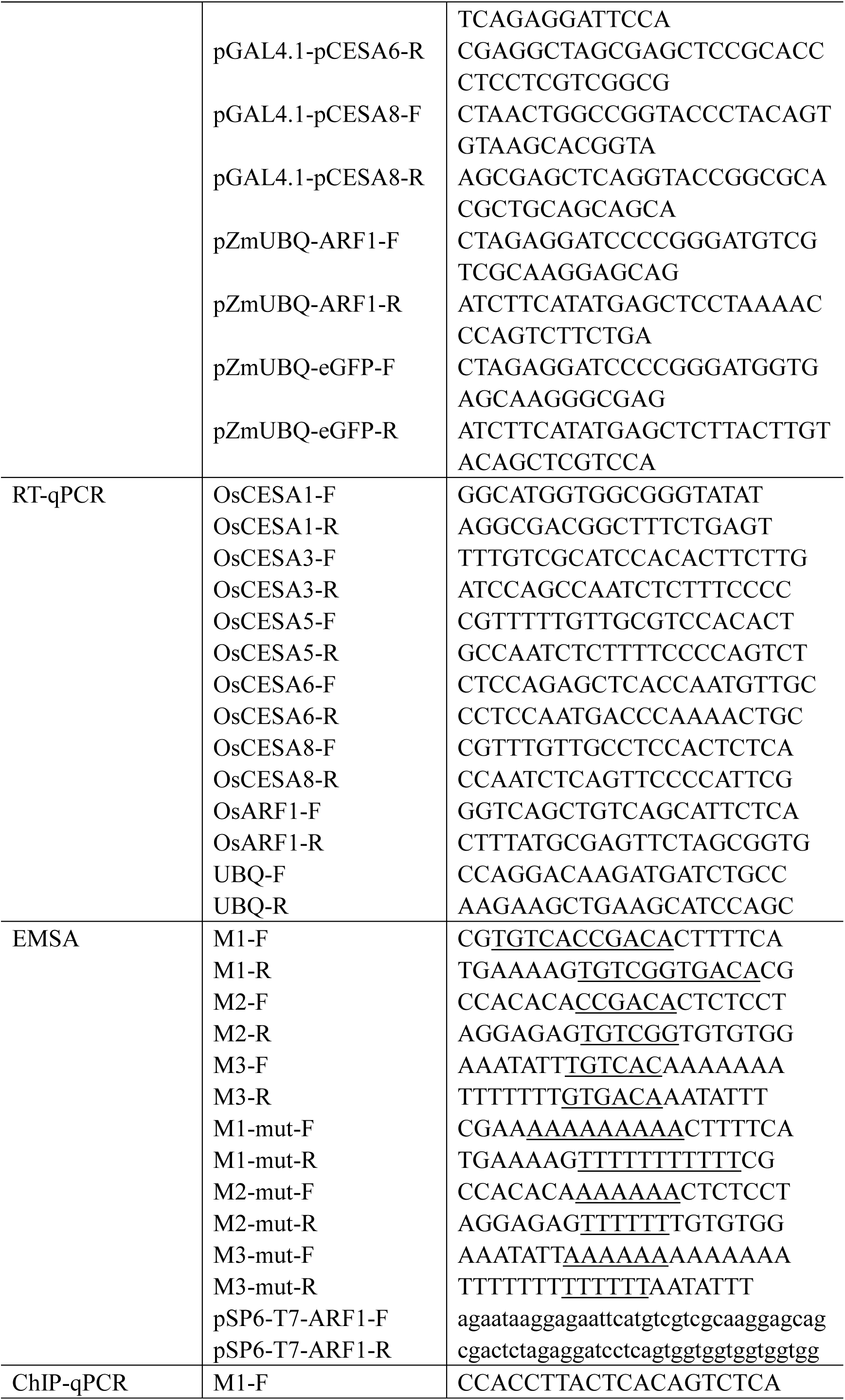
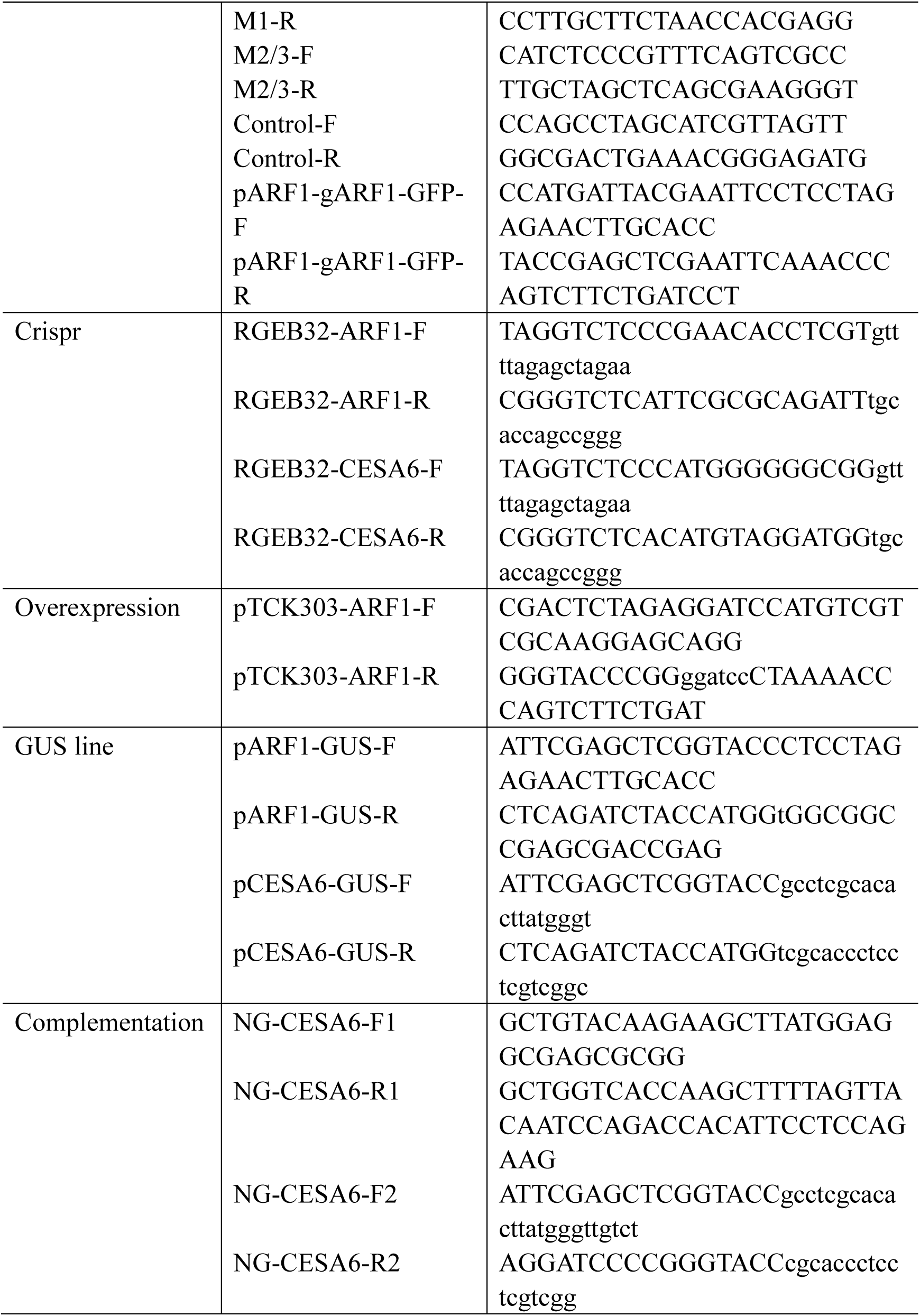
Primers used in this study.

